# The Virtual Lab: AI Agents Design New SARS-CoV-2 Nanobodies with Experimental Validation

**DOI:** 10.1101/2024.11.11.623004

**Authors:** Kyle Swanson, Wesley Wu, Nash L. Bulaong, John E. Pak, James Zou

**Affiliations:** Department of Computer Science, Stanford University; Chan Zuckerberg Biohub - San Francisco; Department of Biomedical Data Science, Stanford University

## Abstract

Science frequently benefits from teams of interdisciplinary researchers. However, most scientists don’t have access to experts from multiple fields. Fortunately, large language models (LLMs) have recently shown an impressive ability to aid researchers across diverse domains by answering scientific questions. Here, we expand the capabilities of LLMs for science by introducing the Virtual Lab, an AI-human research collaboration to perform sophisticated, interdisciplinary science research. The Virtual Lab consists of an LLM principal investigator agent guiding a team of LLM agents with different scientific backgrounds (e.g., a chemist agent, a computer scientist agent, a critic agent), with a human researcher providing high-level feedback. We design the Virtual Lab to conduct scientific research through a series of team meetings, where all the agents discuss a scientific agenda, and individual meetings, where an agent accomplishes a specific task. We demonstrate the power of the Virtual Lab by applying it to design nanobody binders to recent variants of SARS-CoV-2, which is a challenging, open-ended research problem that requires reasoning across diverse fields from biology to computer science. The Virtual Lab creates a novel computational nanobody design pipeline that incorporates ESM, AlphaFold-Multimer, and Rosetta and designs 92 new nanobodies. Experimental validation of those designs reveals a range of functional nanobodies with promising binding profiles across SARS-CoV-2 variants. In particular, two new nanobodies exhibit improved binding to the recent JN.1 or KP.3 variants of SARS-CoV-2 while maintaining strong binding to the ancestral viral spike protein, suggesting exciting candidates for further investigation. This demonstrates the ability of the Virtual Lab to rapidly make impactful, real-world scientific discovery.

## Introduction

Interdisciplinary science research is complex, requiring increasingly large teams of researchers with expertise in diverse fields of science^1–3^. For example, the paper by Jumper et al.^4^ that introduced AlphaFold 2 and later led to the 2024 Nobel Prize in Chemistry included 34 researchers with expertise across computer science, machine learning, bioinformatics, and structural biology. Building and coordinating large teams of researchers who speak different scientific languages and have different scientific priorities is challenging^5,6^. Furthermore, it can be harder for under-resourced groups without connections to many experts across fields to engage in complex, interdisciplinary science, especially when dedicated interdisciplinary research funding is lacking^7^.

One source of broad scientific knowledge and insights that researchers are now turning to is large language models (LLMs), such as ChatGPT^8^ and Claude^9^. These LLMs have been trained on vast quantities of text data, including scientific literature, and they are therefore able to aid researchers in several ways such as by answering science questions, summarizing scientific papers, and writing scientific code^10^. Several studies have explored the scientific capabilities of LLMs by measuring their ability to answer scientific questions, and LLMs have shown high accuracy and can even match or outperform human scientists at these tasks^11–16^.

However, answering individual science questions is very different from engaging in sophisticated research that involves multi-step reasoning across disparate scientific fields with many unknowns. While some prior work has explored the application of LLMs to research, these studies have often focused on a single scientific domain and have explored a relatively narrow set of research questions. For example, ChemCrow is a framework that gives GPT-4 access to chemistry tools and can thus solve components of a chemistry research problem, but it cannot tackle an open-ended, interdisciplinary research problem^17^. Another framework called Coscientist includes GPT-4-powered modules such as a planner and a web searcher to handle several aspects of research^18^. However, Coscientist is primarily applied to relatively standard chemistry tasks such as chemical synthesis planning as opposed to high-level research design across disciplines. In contrast, the AI Scientist aims to use LLMs to perform the entire scientific process from generating a hypothesis to writing code to drafting a paper, but the applications are limited to narrow subfields of machine learning without real-world experiments or validation^19^. Si et al.^20^ similarly explore the use of LLMs for research idea generation and demonstrate promising results when comparing LLM research ideas to human research ideas, but the applications are limited to the field of natural language processing and do not include any implementation of the research ideas.

Here, we introduce the Virtual Lab to overcome these shortcomings via an AI-human research collaboration that performs interdisciplinary science to investigate broad, complex research questions. In the Virtual Lab, a human researcher guides a set of interdisciplinary AI agents^21,22^, such as a biologist or computer scientist, through a set of research meetings that tackle the different phases of a research project. The AI agents are run by an LLM that powers their scientific reasoning abilities with instructions that guide each agent’s scientific expertise and interaction with the other agents and the human researcher. The Virtual Lab architecture is versatile and can thus be applied to a wide variety of interdisciplinary science research projects.

To demonstrate the abilities of the Virtual Lab, we employ the Virtual Lab to tackle a high-impact, real-world, open-ended scientific problem: designing new nanobodies that exhibit binding to the latest variant of SARS-CoV-2. There are myriad ways in which scientists could attempt to design such nanobodies, so the Virtual Lab must reason across multiple subfields of biology and computer science to make a series of interrelated decisions about how best to design these nanobodies. Through a series of meetings, the Virtual Lab develops a novel computational nanobody design workflow that incorporates the protein language model ESM^23^, the protein folding model AlphaFold-Multimer^24^, and the computational biology software Rosetta^25^ to mutate existing nanobodies that bind to the receptor binding domain (RBD) of the spike protein of the original (Wuhan) strain of SARS-CoV-2 into nanobodies that bind to the latest variants of the virus, where an effective binder is lacking^26^. We experimentally validated 92 mutant nanobodies designed by the Virtual Lab and found that over 90% of the nanobodies were expressed and soluble, with two promising candidates showing unique binding profiles to the recent JN.1 and KP.3 spike RBD variants. This result demonstrates the capability of the Virtual Lab’s AI-human collaboration to perform complex, interdisciplinary science research that translates to validated results in the real world.

## Virtual Lab architecture

### Overview

We created the Virtual Lab as a collaboration between a human researcher and a team of large language model (LLM) agents to conduct sophisticated, interdisciplinary research (Fig. 1). The human researcher provides high-level guidance for the LLM agents while the LLM agents both decide on general research directions and design solutions to specific research problems. Each agent is implemented by providing the underlying LLM (e.g., GPT-4o) with specific roles (e.g., a biologist agent) and domain-specific tools (e.g., AlphaFold). The Virtual Lab performs research in two primary ways: team meetings and individual meetings. In both cases, the human researcher provides an initial agenda to guide the discussion, and then the agents discuss how to address the agenda. In team meetings, all of the agents discuss a broad research question and work together to come up with an answer. In individual meetings, a single agent is given a more specific task to accomplish, such as writing code for a machine learning model, and the agent either works alone or in conjunction with another agent that provides critical feedback.

**Fig. 1.**
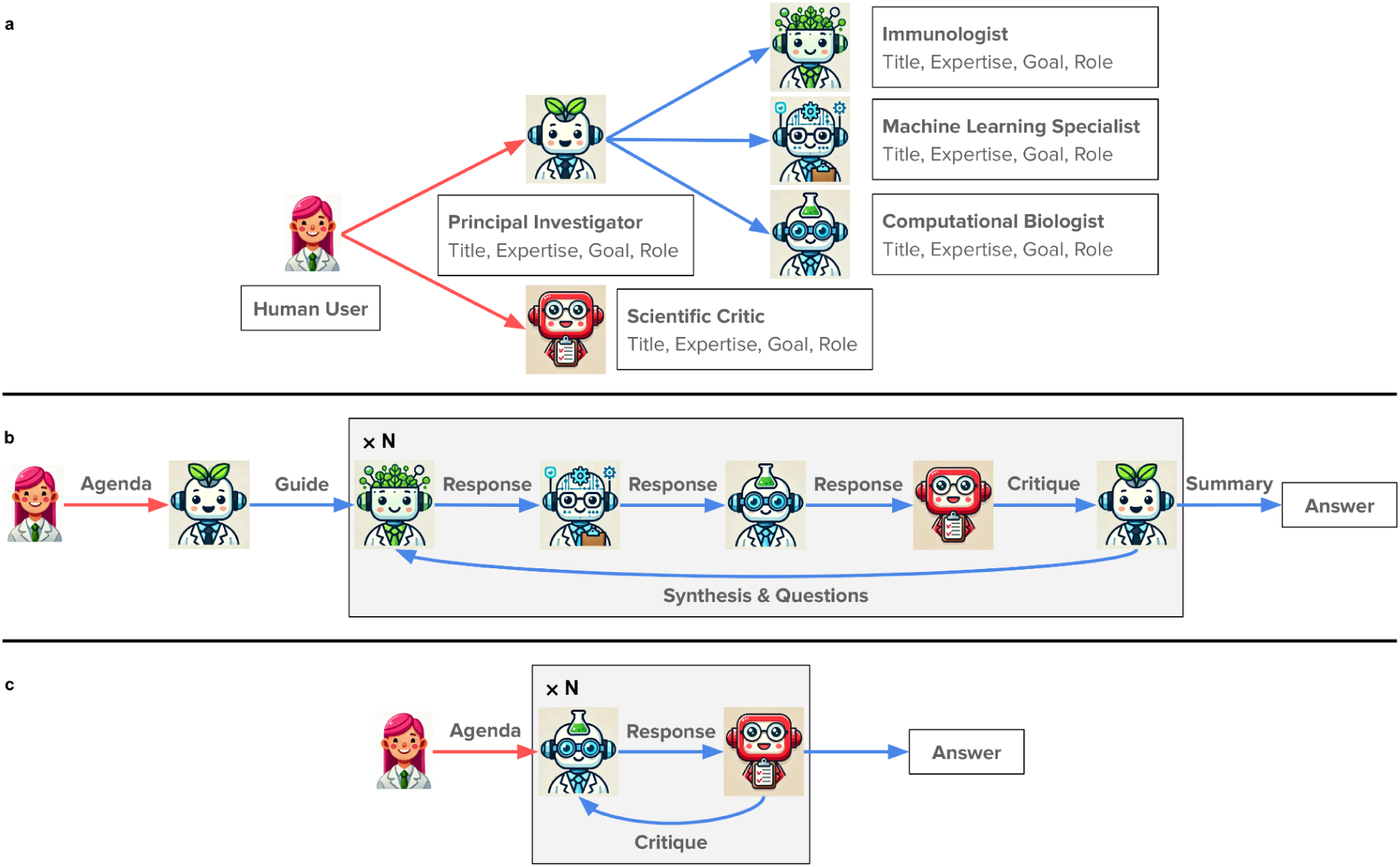
Virtual Lab architecture. **a**, The workflow for designing agents in the Virtual Lab. Each agent is specified with four criteria: title, expertise, goal, and role. The human researcher in the Virtual Lab specifies these criteria to define the Principal Investigator (PI) agent and the Scientific Critic agent. Then, given a short description of the project by the human researcher, the PI agent automatically creates several scientist agents to work on the project by specifying their title, expertise, goal, and role, using its own prompt as an example. **b**, The workflow for a team meeting in the Virtual Lab. The human researcher writes an agenda for the meeting specifying the topic of discussion. The PI agent begins the meeting by providing initial thoughts and agenda questions as a guide for the remaining agents. Then, over the course of N rounds of discussion, each scientist agent provides its response, followed by a critique by the Scientific Critic agent, with the PI agent then synthesizing the discussion and asking follow-up questions. Finally, after the N rounds of discussion, the PI agent summarizes the discussion and provides an answer regarding the meeting agenda. **c**, The workflow for an individual meeting. The human researcher writes an agenda for the meeting specifying the topic of discussion. Then, the scientist agent tasked with the individual meeting provides a response to the agenda, which is critiqued by the Scientific Critic. In each round, the scientist agent improves its answer based on feedback from the Scientific Critic. Finally, after the N rounds, the scientist agent provides its final, improved answer.

Through a series of team meetings and individual meetings, the Virtual Lab tackles a complex research project.

### Agents

Each LLM agent in the Virtual Lab is defined with a prompt that specifies four key criteria (Fig. 1a).

1. **Title:** The name of the agent.
2. **Expertise:** The scientific expertise the agent has.
3. **Goal:** The ultimate goal of the agent in the context of the research project.
4. **Role:** The specific role that the agent will play in the research project.

The agents of the Virtual Lab are led by an agent called the Principal Investigator (PI). The PI agent has expertise in artificial intelligence for scientific research with a goal of maximizing the scientific impact of research and with the role of guiding the research project (full prompt in the Appendix). The PI agent then automatically creates a set of scientist agents (e.g., a biologist or a computer scientist) that are appropriate for the research project based on a short description of the project written by the human researcher. The PI defines these scientist agents by specifying each agent’s title, expertise, goal, and role, using its own prompt as an example.

In addition to the PI and scientist agents, we find it useful to create an explicit critic agent to catch errors and oversights from the other agents and to give critical feedback on answers provided by the other agents^27^. Therefore, a Scientific Critic agent (prompt in the Appendix) can be added to any team meeting or individual meeting to provide critical feedback to the other agents.

### Meetings

Interactions in the Virtual Lab happen through meetings, which can either be *team meetings* with all the agents or *individual meetings* with a single agent (and optionally the critic agent). Both types of meetings share the following set of inputs that structure the meeting.

1. **Agenda:** (required) A description of the scientific topic to be discussed during the meeting.
2. **Agenda questions:** (optional) A set of questions that the agents must answer by the end of the meeting.
3. **Agenda rules:** (optional) A set of rules that the agents must follow during the meeting.
4. **Summaries:** (optional) Agent-written summaries of previous meetings to provide information about previous decisions.
5. **Contexts:** (optional) Additional information (e.g., scientific papers) for the agents to take into consideration.
6. **Rounds:** (required) The number of rounds (typically N=3) of discussion among the agents.

The team and individual meetings differ in terms of the agents that participate in the meeting and the prompts that guide the flow of the meeting.

### Team meeting

In team meetings, all the agents (PI agent, scientist agents, and Scientific Critic agent) participate in a conversation to address a broad research topic (Fig. 1b). First, the human researcher writes an agenda for the team meeting along with any applicable agenda questions and agenda rules. The team meeting then begins with an automatically constructed prompt (see Appendix) then introduces the agents, agenda, agenda questions (if any), and agenda rules (if any) and describes the flow of the meeting, which involves multiple rounds of discussion. The PI agent is prompted to start the discussion by providing their initial thoughts and any guiding questions that they want to ask the team. Then, each scientist agent and the Scientific Critic agent are prompted one-by-one (in an order set by the human researcher) to provide their thoughts on the ongoing discussion given everything that has been said by the other agents. At the end of a round of discussion, the PI agent synthesizes the points raised by each agent, makes decisions based on agent input, and asks follow-up questions to further the discussion. After N rounds of discussion (with N set by the human researcher), the PI agent summarizes the discussion for future meetings, provides a recommendation regarding the agenda, and answers the agenda questions (if any). The human researcher in the Virtual Lab can then read just this final response by the PI agent, thus benefiting from the extensive discussions among the LLM agents while only needing to read the final short response to understand the decisions that were made.

### Individual meeting

In individual meetings, a single agent tackles a specific task that falls within their area of expertise, optionally with critical feedback provided by the Scientific Critic agent (Fig. 1c). To start an individual meeting, the human researcher in the Virtual Lab selects the agent that will participate. An automatically constructed prompt (see Appendix) introduces the agenda, agenda questions (if any), and agenda rules (if any) and then immediately asks the agent for a response. If the individual meeting has zero rounds (N=0), then the agent provides a response and the meeting ends. If the individual meeting includes one or more rounds (N≥1), then in each round, the agent provides a response and then the Scientific Critic agent provides critical feedback to improve the agent’s response. After these rounds, the selected agent responds one more time to provide the final, improved answer.

### Parallel meetings

To improve the expected quality and comprehensiveness of answers for a given meeting, the same meeting (same agents, same prompts) can be run multiple times in parallel to produce multiple answers (Supplementary Fig. 1). Then, an individual meeting with the appropriate agent (i.e., the PI agent for team meetings or the relevant scientist agent for individual meetings) is run to merge the summaries of each of the parallel meetings into a single answer that incorporates the best elements from each of the parallel meetings. To boost creativity while producing a consistently high-quality answer, each of the parallel meetings is run with a higher “creative” temperature of 0.8 while the single merge meeting is run with a lower “consistent” temperature of 0.2^28^. Parallel meetings are similar in nature to the method of majority voting from multiple LLM queries^29^, but the Virtual Lab’s parallel meetings use a more complex and flexible merging of answers via a meeting with an LLM agent.

## Virtual Lab for nanobody design

### Overview

Given the flexibility of the Virtual Lab architecture, the Virtual Lab can be applied to a wide variety of interdisciplinary research projects by adapting the agents and the flow of team and individual meetings to the specific project’s goals and constraints. As a demonstration in the domain of biological research, we applied the Virtual Lab with GPT-4o^30^ powering the agents to design antibodies or nanobodies that can bind to the spike protein of the KP.3 variant of SARS-CoV-2, which was one of the latest variants at the time of this work^26^ (Fig. 2). This is an important and challenging problem because SARS-CoV-2 is rapidly evolving resistance to existing antibody/nanobody therapies, so quickly developing new antibody/nanobody therapies that overcome this resistance and bind to the latest variants is crucial to treating those who are infected^31,32^. The Virtual Lab tackles this problem by rapidly creating a computational workflow to design antibodies or nanobodies for the KP.3 variant of SARS-CoV-2, which can then be experimentally validated by human biologists. The Virtual Lab created the computational antibody/nanobody design process in five phases.

1. **Team selection:** An individual meeting with the PI to define a set of scientist agents to work on the project.
2. **Project specification:** A team meeting to specify the project direction by deciding on key high-level details.
3. **Tools selection:** A team meeting to brainstorm machine learning and/or computational tools for nanobody design.
4. **Tools implementation:** A series of individual meetings with different scientist agents to implement three components of the nanobody design workflow—ESM, AlphaFold-Multimer, and Rosetta.
5. **Workflow design:** An individual meeting with the PI to determine the workflow for applying these computational tools.

**Fig. 2.**
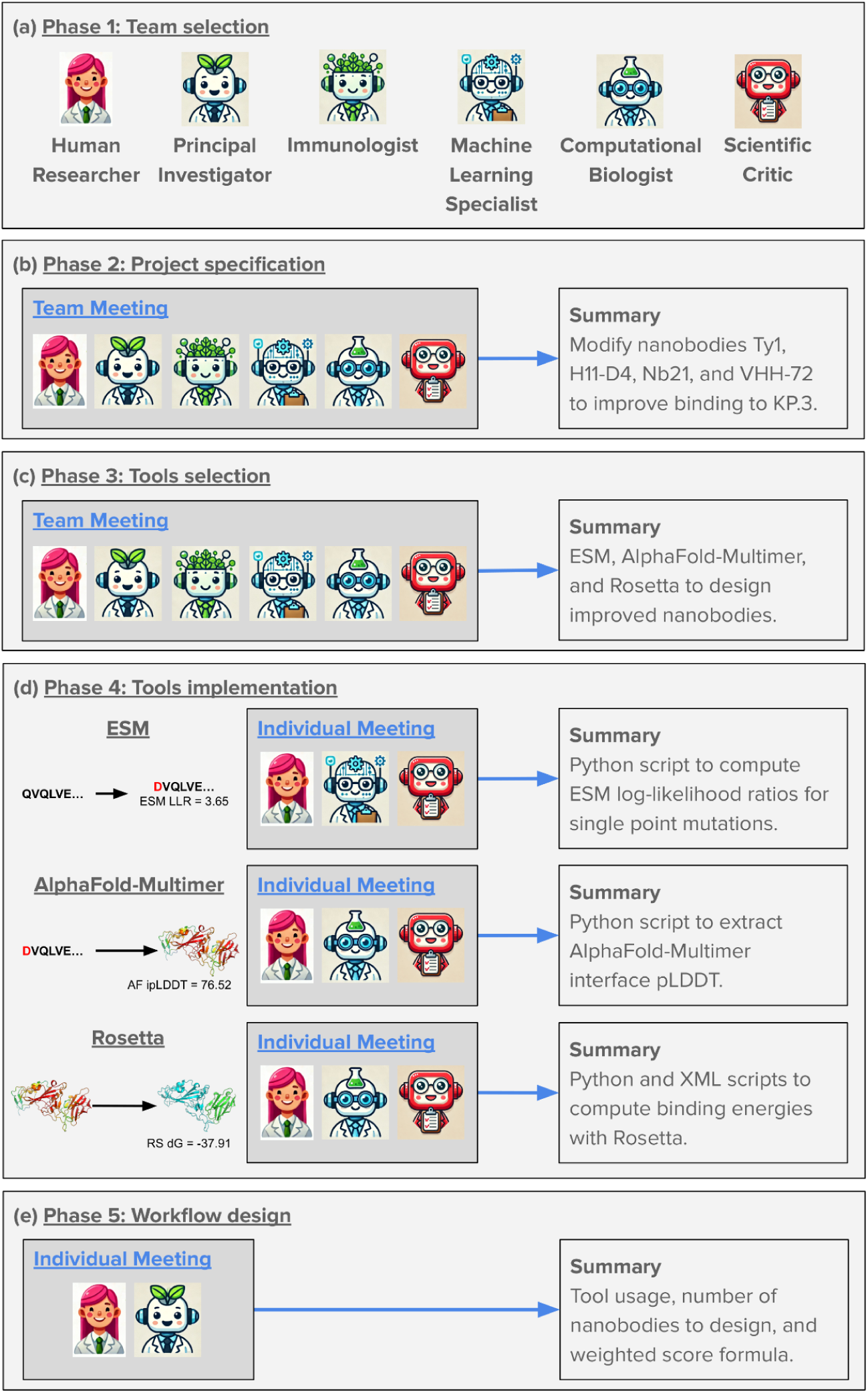
Virtual Lab for nanobody design. The workflow used to apply the Virtual Lab to nanobody design for the latest variant of SARS-CoV-2. **a**, The workflow begins with the human researcher defining the Principal Investigator (PI) and Scientific Critic agents by specifying their title, expertise, goal, and role. Then, the PI agent creates a team of three scientist agents for the project. **b**, A team meeting discusses the project specification, and the agents make decisions such as whether to design antibodies or nanobodies. **c**, In another team meeting, the agents suggest a set of computational tools for nanobody design, including ESM, AlphaFold-Multimer, and Rosetta. **d**, In a series of individual meetings, the Machine Learning Specialist and Computational Biologist, with helpful feedback from the Scientific Critic, write code and subsequently improve that code for the ESM, AlphaFold-Multimer, and Rosetta components of the nanobody design workflow. **e**, In an individual meeting, the PI agent decides the workflow for using the three computational tools to design and select mutated nanobody candidates.

These phases are discussed in more detail below, and complete prompts for each phase are in the Appendix.

### Team selection

First, Virtual Lab used an individual meeting with the PI agent (run five times in parallel followed by a merge meeting) to create a set of scientist agents for the project. The meeting agenda contained a background prompt about antibody/nanobody design for the spike protein of the recent KP.3 SARS-CoV-2 variant and a request for the PI agent to select a team of three scientist agents for the project using the same agent structure (title, expertise, goal, role) as the PI’s own definition. The PI decided to create an Immunologist, a Machine Learning Specialist, and a Computational Biologist.

### Project specification

Next, in a team meeting, the full team of agents (PI, Immunologist, Machine Learning Specialist, Computational Biologist, and Scientific Critic) discussed some of the specifics of the project beyond the general background prompt. This meeting’s agenda asked the agents to consider design choices such as whether to pursue antibodies or nanobodies and whether to modify existing antibodies/nanobodies or design new ones *de novo*. This team meeting was run in five parallel iterations followed by an individual merge meeting with the PI agent and the Scientific Critic to produce the best answer. Below are some of the PI’s final merged answers to the agenda questions.

#### 1. Will you design standard antibodies or nanobodies?

**Agent Answer:** Nanobodies

**Agent Justification:** Nanobodies offer superior stability, tissue penetration, and ease of production, making them ideal for targeting conserved and cryptic epitopes on the SARS-CoV-2 spike protein. This choice is consistently supported across all summaries, emphasizing their advantages in accessing unique epitopes and potential for broad-spectrum activity.

#### 2. Will you design antibodies/nanobodies de novo or will you modify existing antibodies/nanobodies (choose only one)?

**Agent Answer:** Modify existing nanobodies

**Agent Justification:** Modifying existing nanobodies allows us to leverage established data, providing a quicker and more reliable path to developing broad-spectrum candidates. This approach is supported by the availability of structural data and known efficacy, which can accelerate the optimization process. This decision is consistently recommended across all summaries, highlighting the efficiency and feasibility of building on known frameworks.

#### 3. If modifying existing antibodies/nanobodies, which precise antibodies/nanobodies will you modify (please list 3-4)?

**Agent Answer:** Ty1, H11-D4, Nb21, VHH-72

**Agent Justification:** These candidates have demonstrated strong neutralizing capabilities against SARS-CoV-2 and have available structural data, making them excellent candidates for modification. Ty1 and H11-D4, for example, have shown efficacy in neutralizing multiple variants, while Nb21 and VHH-72 offer unique binding properties that can be optimized for broader cross-reactivity. Literature supports their efficacy, with studies indicating their ability to bind conserved epitopes and maintain stability under various conditions.

Given these decisions, the following phases proceeded with nanobody design by modifying the four nanobodies suggested by the Virtual Lab (Ty1^33^, H11-D4^34^, Nb21^35^, and VHH-72^36^), which are specific to the ancestral Wuhan spike protein, to increase their affinity to the spike protein of the KP.3 variant of SARS-CoV-2. Furthermore, the Virtual Lab suggested prioritizing “enhancing interactions with the receptor-binding domain of the spike protein by altering residues that contribute to binding affinity,” so the Virtual Lab subsequently focused on developing nanobodies that bind to the receptor binding domain (RBD) of the KP.3 spike protein.

### Tools selection

After specifying the project direction, the Virtual Lab next needed to pick a set of computational tools to modify the selected nanobodies. To accomplish this, the Virtual Lab ran a team meeting asking the agents to list several machine learning and/or computational tools that could be used for nanobody design, with emphasis on pre-trained models for simplicity. Similar to the project selection meeting, this team meeting was run with five parallel iterations followed by a merge meeting with the PI and Scientific Critic. The agents decided to use ESM^23^, AlphaFold-Multimer^24^, and Rosetta^25^ as the components of its computational nanobody design workflow.

### Tools implementation

With the project well-specified and a set of computational nanobody tools chosen, the Virtual Lab then worked on implementing these tools for nanobody design. For each tool, the Virtual Lab selected the most appropriate scientist agent via an individual meeting with the PI. Then for each tool, the Virtual Lab ran an individual meeting with the selected scientist agent and the Scientific Critic (five parallel meetings followed by a merge meeting run by the scientist agent) to implement the tool. These meetings included a set of agenda rules that specify how code should be written, e.g., with good documentation and without leaving functions undefined. These initial implementations contained small errors that needed correction, so the Virtual Lab then ran a single follow-up individual meeting (no parallelization or Scientific Critic) with the scientist agent to automatically fix all the errors that arose (see Appendix).

### ESM usage

The Machine Learning Specialist agent was responsible for writing a Python script to identify the most promising point mutations to a nanobody sequence based on the ESM log-likelihood ratio (LLR) of the mutant sequence compared to the input sequence. The agent wrote a 130-line Python script with three functions: a main function to run the script, a function to parse command-line arguments (e.g., the input nanobody sequence), and a function that uses a pre-trained ESM model to compute log-likelihood ratios for point mutations.

Notably, the Virtual Lab’s ESM nanobody mutation analysis is similar to the ESM-based antibody design process of Hie et al.^37^, but it differs in important ways. The Virtual Lab computes the ESM LLR of mutant sequence *x*’ with mutation at position *i* compared to the input sequence *x* as 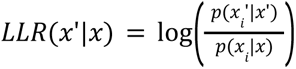, where 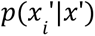 is the ESM probability of the mutant amino acid 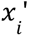 at position *i* in the mutant sequence *x*’ and *p*(*x_i_*|*x*) is the ESM probability of the original amino acid *x_i_* at position *i* in the input sequence *x*. In contrast, Hie et al. use 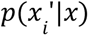 as the numerator in the ratio instead of 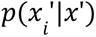, thereby computing the probability of the mutant amino acid in the *input* sequence rather than the probability of the mutant amino acid in the *mutant* sequence. Additionally, the Virtual Lab only uses the ESM-1b^38^ model while Hie et al. use a consensus of ESM-1b and five ESM-1v^39^ models.

### AlphaFold-Multimer usage

To use AlphaFold-Multimer, the Virtual Lab asked the Computational Biologist agent to write a Python script that processes a predicted nanobody-spike complex structure from AlphaFold-Multimer and outputs the interface pLDDT (ipLDDT), which is a measure of the confidence of the binding interface between the nanobody and the spike protein that has previously been shown to correlate with antibody-antigen binding affinity^40^. Computing the ipLDDT values across multiple proposed nanobodies requires reading a PDB file for each predicted nanobody-spike complex and writing as output a single CSV file with the ipLDDT from every complex. The Computational Biologist wrote a 144-line Python script with five functions: a main function to run the whole script, a function to check whether a PDB file contains a protein structure in the correct format, a function to identify the residues in the interface between the two proteins, a function to calculate the ipLDDT, and a function to run the ipLDDT calculation on every PDB file in a directory and save the results to a single CSV file.

### Rosetta usage

The Computational Biologist was also responsible for using Rosetta to calculate nanobody-spike binding energies as a metric for measuring the quality of each mutated nanobody. Given a PDB file with a predicted nanobody-spike structure from AlphaFold-Multimer, the Computational Biologist was asked to write a RosettaScripts XML file to load the PDB file, calculate the binding energy, and save the binding energy to a Rosetta score file. Additionally, the agent was asked to write a Python script that loads all the score files in a directory and saves a CSV file with the binding energy of every nanobody-spike complex.

The Computational Biologist wrote a 30-line RosettaScripts XML file that first relaxes the nanobody-spike structure and then computes the binding energy (dG-separated in Rosetta terminology) of the interface using the REF15 scoring function. The Computational Biologist then wrote a 71-line Python script with two functions: a main function to run the whole script and a function to extract the binding energy score from a given Rosetta score file.

### Workflow design

Finally, the Virtual Lab ran an individual meeting with the PI agent to design a workflow that uses ESM, AlphaFold-Multimer, and Rosetta to design nanobodies. For each of the four starting nanobody candidates, the PI agent decided to run ESM to evaluate all possible point mutations and then to select the top 20 mutations by ESM log-likelihood ratio. Each of these 20 mutant sequences would then be evaluated by both AlphaFold-Multimer and Rosetta. These 20 sequences would then be ranked and the top five would be selected using the following weighted score designed by the PI agent:

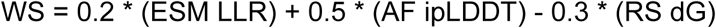

where WS is the weighted score, ESM LLR is the ESM log-likelihood ratio between the mutated sequence and the input sequence, AF ipLDDT is the AlphaFold-Multimer ipLDDT binding interface confidence, and RS dG is the Rosetta dG-separated binding energy value. The PI correctly uses a negative weight for the Rosetta value since a more negative binding energy is better. The top five sequences according to WS then serve as the starting sequences for the next round of mutation, with 4 rounds of mutation in total depending on time constraints and improvements in the WS across rounds.

## Results

### Nanobody design

The Virtual Lab ran the nanobody design computational workflow (Fig. 3a) to design improved nanobody candidates for the KP.3 variant of SARS-CoV-2, which was one of the latest variants at the time of this work. The workflow was run independently for each of the four nanobodies suggested by the agents: Ty1, H11-D4, Nb21, and VHH-72. Below, we describe the workflow in terms of a single starting nanobody for simplicity.

**Fig. 3.**
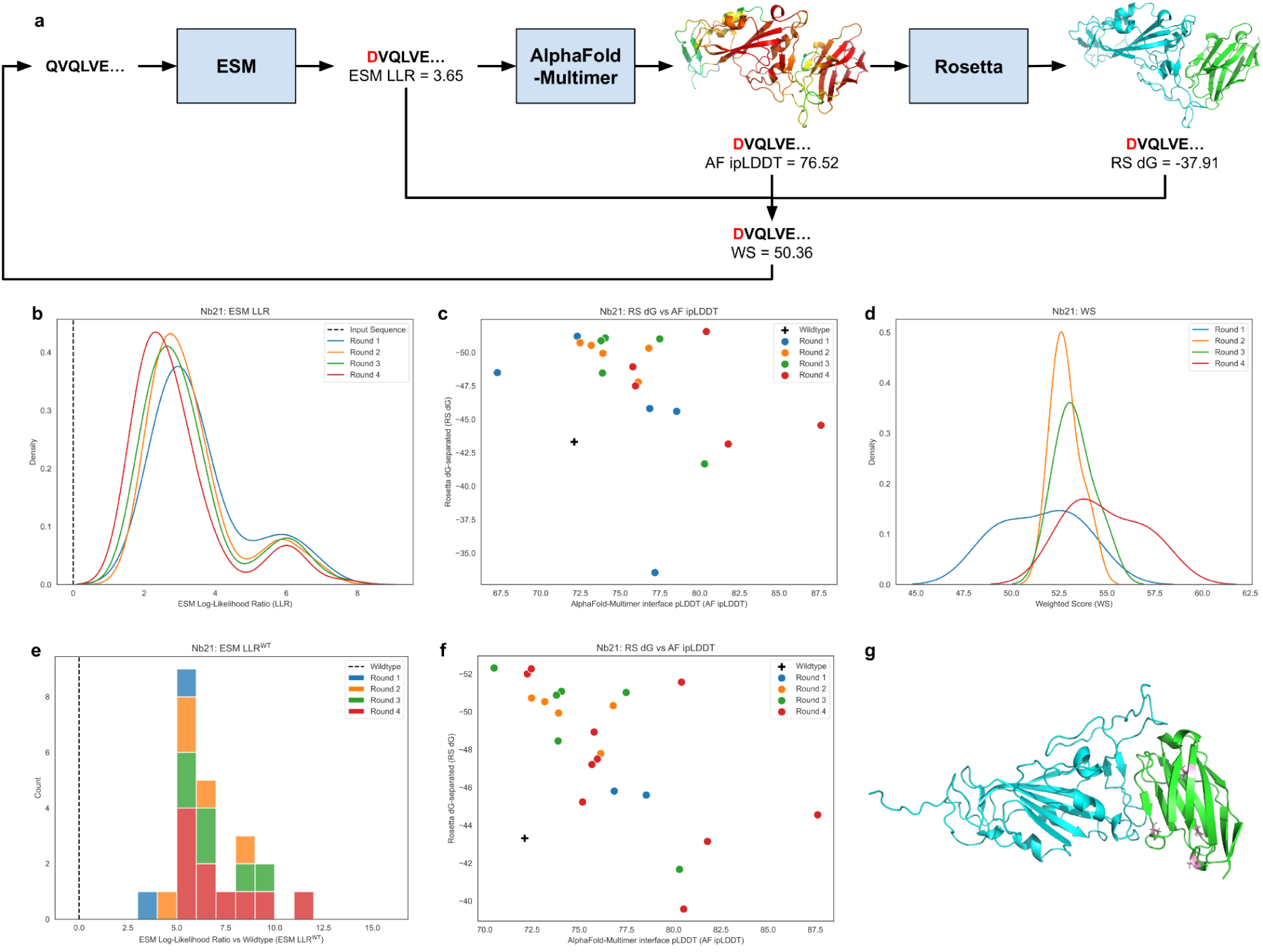
Nb21 nanobody analysis. **a**, Each round of nanobody design begins with a nanobody sequence. ESM computes the log-likelihood ratio (ESM LLR) of every single point mutation to the input sequence. The top 20 mutant sequences by ESM LLR are run through AlphaFold-Multimer to predict the structure of the complex of nanobody and SARS-CoV-2 spike protein, and the interface pLDDT (AF ipLDDT) confidence value is computed. This complex is then passed to Rosetta, which relaxes the structure and computes a binding energy (RS dG). These three scores are combined into a weighted score (WS) that is used to select the top five mutant nanobodies for the next round of optimization. **b-d**, Evolution of mutant nanobody scores across four rounds of optimization, with improvements in ESM LLR, AF ipLDDT, and RS dG across the rounds. **b**, The distribution of ESM LLR values for proposed Nb21 mutant nanobodies across each round of optimization, with ESM LLR values computed relative to the input nanobody sequence from the previous round. Shown are the ESM LLR values of the top 20 proposed mutant nanobodies per input nanobody, i.e., the proposed mutant nanobodies that proceed to AlphaFold-Multimer and Rosetta analysis. **c**, The AF ipLDDT and the RS dG of the top five proposed nanobodies, selected by WS, at the end of each round of optimization. **d**, The distribution of WS values of the top five proposed nanobodies at the end of each round of optimization. **e-g**, Analysis of the final set of 23 mutant nanobodies selected across all rounds of optimization, which exhibit superior ESM LRR^WT^, AF ipLDDT, and RS dG values compared to the wild-type nanobody. **e**, The distribution of ESM LLR^WT^ values (ESM LLR of the mutant sequence compared to the wild-type sequence) for the selected nanobodies and the wild-type nanobody. **f**, The AF ipLDDT and RS dG values of the selected nanobodies and the wild-type nanobody. **g**, The structure (predicted by AlphaFold-Multimer followed by Rosetta relaxation) of the receptor binding domain of the spike protein of the KP.3 variant of SARS-CoV-2 (cyan) and the proposed nanobody mutant Nb21 I77V-L59E-Q87A-R37Q (green). Side chains are shown for all residues within 4Å of the opposite chain (i.e., interface residues). Mutant residues in the proposed nanobody are shown in pink.

The Virtual Lab workflow began with “round 0,” which evaluated the wild-type nanobody sequence without introducing any mutations. ESM LLR was assigned to zero since the wild-type nanobody sequence was unmodified. Then, the Virtual Lab ran AlphaFold-Multimer (via LocalColabFold^41^ version 1.5.5) on the nanobody sequence and the sequence of the receptor binding domain (RBD) of the KP.3 spike protein to produce a predicted structure of the complex. Next, the Virtual Lab computed the AF ipLDDT as a measure of confidence in the binding interface of the complex. Then, the Virtual Lab ran Rosetta to relax the complex and compute the RS dG value as an estimate of the binding energy. Finally, the Virtual Lab computed the weighted score (WS) of the wild-type nanobody.

In round 1, the Virtual Lab ran ESM to calculate the ESM LLR of every possible single point mutation to the wild-type nanobody. The top 20 mutated sequences by ESM LRR were retained. For each of these 20 mutated sequences, AlphaFold-Multimer and Rosetta were applied in the same way as for the wild-type sequence. The Virtual Lab then computed the WS for each of the 20 mutated sequences and selected the top five sequences for the next round. In rounds 2-4, the Virtual Lab applied the same procedure but now starting with five input sequences to the ESM LLR script, resulting in 100 top mutated sequences (20 proposed mutant sequences for each of the five input sequences). These sequences were analyzed by AlphaFold-Multimer and Rosetta, and the top five of these 100 sequences were selected at the end of each round by their WS.

After running all four rounds of mutation, the Virtual Lab needed to select the best mutated nanobody sequences across all four rounds for experimental validation. Doing so required using a slight variant of the weighted score (WS). In each round, the WS used the ESM LLR calculated as a ratio between the proposed mutant sequence and the input sequence for that round (i.e., an output sequence from the previous round), which differ by a single mutation.

However, in order to fairly select the best sequences across different rounds with different numbers of mutations, an alternate ESM log-likelihood ratio, which we call the ESM LLR^WT^, was computed between each proposed mutant sequence (with one to four mutations) and the wild-type sequence. The Virtual Lab then scored all mutant nanobody sequences using the WS^WT^, which is the weighted score calculated using the ESM LLR^WT^ in place of the ESM LLR. The top 23 mutant sequences were selected for experimental validation along with the wild-type sequence as a point of reference.

### Computational nanobody analysis

The successive rounds of optimization improved the quality of the proposed mutant nanobody sequences according to the three metrics of ESM LLR, AF ipLDDT, and RS dG. Fig. 3b-g shows relevant metrics for Nb21, while results are similar for Ty1 (Supplementary Fig. 2), H11-D4 (Supplementary Fig. 3), and VHH-72 (Supplementary Fig. 4). Table 1 shows scores for the wild-type sequence and some of the mutant sequences that were selected for experimental validation.

**Table 1.** Nanobody score analysis. The scores of each wild-type nanobody and examples of the mutant nanobodies that were selected for experimental validation. ESM LLR^WT^: ESM log-likelihood ratio between the mutant nanobody sequence and the wild-type sequence. AF ipLDDT: AlphaFold-Multimer interface pLDDT for the nanobody-spike complex. RS dG: Rosetta dG-separated binding energy value. WS^WT^: Weighted score combining ESM LLR^WT^, AF ipLDDT, and RS dG.

| Name | ESM LLR<sup>WT</sup> | AF ipLDDT | RS dG | WS<sup>WT</sup> |
|---|---|---|---|---|
| Ty1 | 0.00 | 71.83 | -41.51 | 48.36 |
| Ty1 V32F-G59D-N54S-F32S | 3.51 | 86.06 | -28.69 | 52.34 |
| H11-D4 | 0.00 | 68.18 | -38.93 | 45.77 |
| H11-D4 A14P-Y88V-K74T-R27L | 10.67 | 84.02 | -35.04 | 54.66 |
| Nb21 | 0.00 | 72.11 | -43.32 | 49.05 |
| Nb21 I77V-L59E-Q87A-R37Q | 7.47 | 80.41 | -51.56 | 57.17 |
| VHH-72 | 0.00 | 66.46 | -20.90 | 39.50 |
| VHH-72 R27Y-E31D-F37V-D89E | 8.82 | 69.51 | -53.76 | 52.65 |

In each round, the top sequences selected by ESM LLR had log-likelihood ratios of roughly 1-8, indicating that each subsequent round of mutation improved the overall quality of the nanobody compared to the input sequence from the previous round (Fig. 3b). This is according to ESM’s internal understanding of nanobody likelihood, which does not take the antigen (spike protein) into account but does understand overall nanobody quality. The top mutant nanobody sequences selected by ESM LLR in each round generally had improved structural complexes with the KP.3 spike protein based on improved AF ipLDDT, improved RS dG, or both (Fig. 3c). The WS values of the top five sequences at the end of each round improved (Fig. 3d), even when using the ESM LLR instead of the ESM LLR^WT^ that corrects for the effect of multiple mutations and not just the most recent mutation.

After correction, the ESM LLR^WT^ for the final selected 23 sequences showed a large improvement over the wild-type sequence (Fig. 3e). These selected sequences also had improved AF ipLDDT and RS dG scores compared to the wild-type (Fig. 3f). An example of the AlphaFold-Multimer predicted structure of a top scoring mutant nanobody is shown in Fig. 3g. Notably, the final set of 23 selected nanobody sequences includes sequences with different numbers of mutations (i.e., from different rounds) and with a different balance of ESM LLR^WT^, AF ipLDDT, and RS dG values, allowing for a diversity of potential improvements to the wild-type nanobody.

Applying this workflow to each of the four starting nanobodies resulted in 92 final selected sequences (23 per starting nanobody). All 92 mutant nanobodies had a positive ESM LLR, indicating that ESM preferred the mutant over the wildtype. Among the 92 mutant nanobodies, 78 (85%) had an AF ipLDDT greater than their respective wildtype nanobody, and 32 (35%) had an AF ipLDDT ≥ 80, which is in line with the AF ipLDDT scores of high accuracy AlphaFold-Multimer antibody-antigen structural models in prior work^40^. Furthermore, 60 (65%) had an RS dG less (better) than their respective wildtype nanobody, and 23 (25%) of the 92 mutants had an RS dG ≤ −50, which is in line with strong Rosetta binding energy values of nanobodies or antibodies in complex with the SARS-CoV-2 receptor binding domain in prior work^25,42^.

### Nanobody experimental validation

To validate the nanobodies designed by the Virtual Lab, we conducted a set of experiments to measure their binding to a panel of spike receptor binding domain (RBD) proteins (Fig. 4). We first overexpressed each nanobody in *E. coli*, followed by isolation of soluble protein from the periplasm. The designed nanobodies show excellent expression, with 38% (35 of 92) of the designs showing high to very high expression levels (Fig. 5a, Supplementary Fig. 5) and only 6.5% (6 of 92) of the designs showing very little to no soluble expression. Thus, the mutations proposed by the Virtual Lab are well tolerated and do not cause large-scale misfolding or aggregation of the nanobodies.

**Fig. 4.**
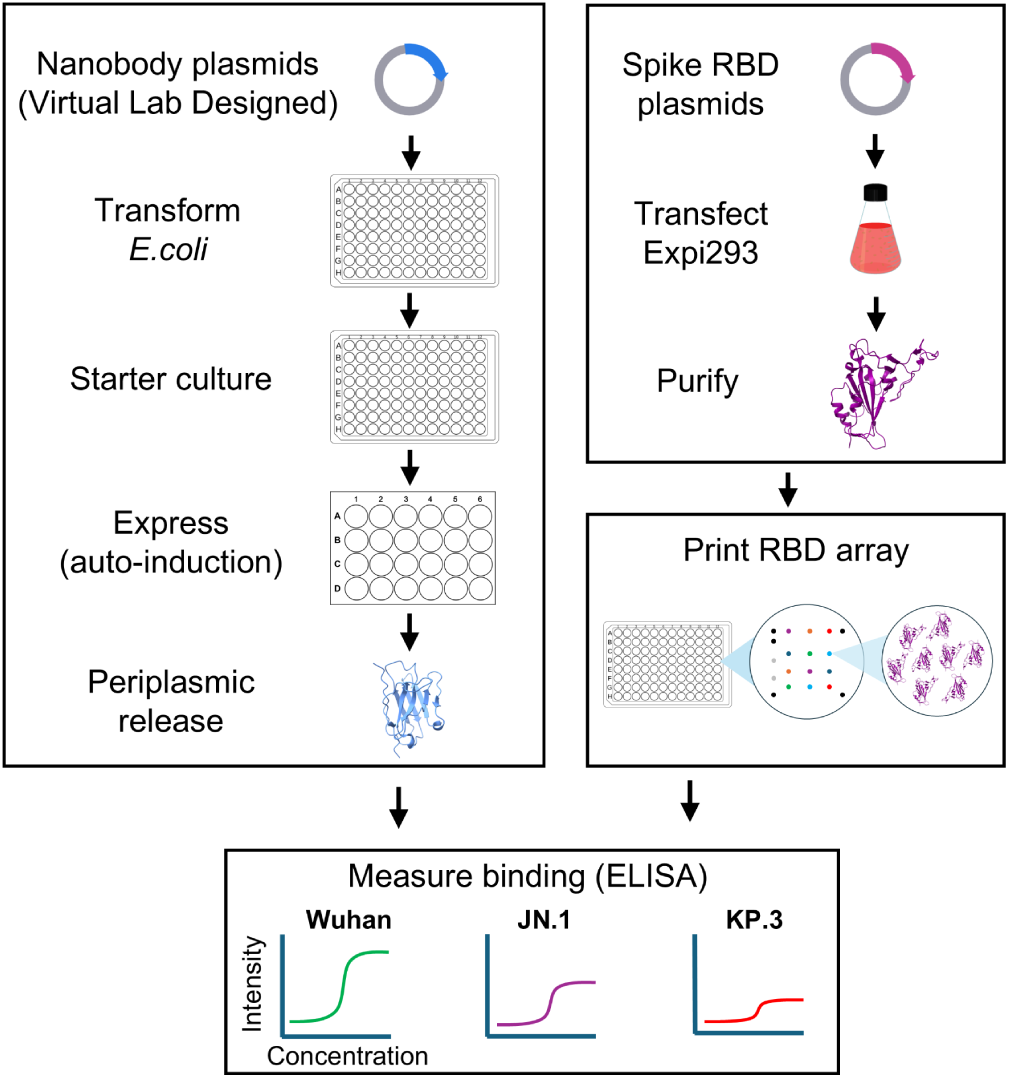
Workflow for nanobody experimental validation. The four categories of experiments (nanobody expression, SARS-CoV-2 spike RBD expression, antigen array printing, and multiplexed ELISA) are enclosed in boxes. The ribbons representation of a nanobody (blue) and the RBD (purple) were rendered with ChimeraX^58^ from PDB accession numbers 6XZN and 6M0J, respectively. Unique RBD and control proteins of the array are shown as colored spots with fiducial markers shown as black spots.

**Fig. 5.**
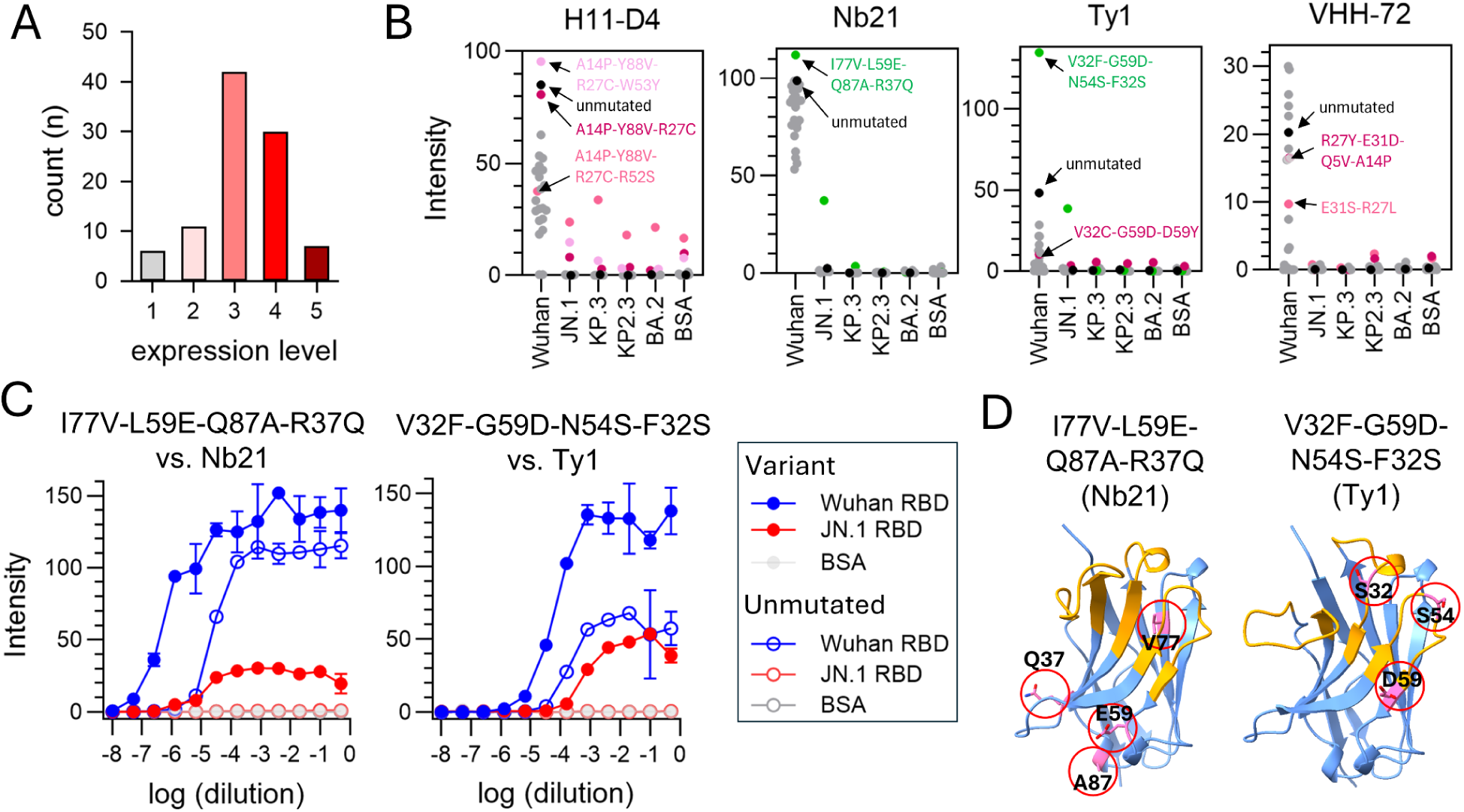
Experimental validation of Virtual Lab nanobodies. **a**, Histogram of nanobody expression levels. A 5-point scale (1 = very low, 2 = low, 3 = average, 4 = high, 5 = very high) was used to classify the amount of nanobody in the soluble periplasmic extract. SDS-PAGE data used for the graph is shown in Supplementary Fig. 5. **b**, ELISA binding profiles of nanobodies to a panel of antigens. For each SARS-CoV-2 RBD protein and BSA (x-axis), individual spots represent the ELISA binding intensity (y-axis) of each nanobody. Unmutated nanobodies (H11-D4, Nb21, Ty1, and VHH-72) are shown in black and nanobodies exhibiting high non-specific binding are shown in shades of red (light pink, pink, and magenta). The Nb21 mutant (I77V-L59E-Q87A-R37Q) and the Ty1 mutant (V32F-G59D-N54S-F32S) that show binding to JN.1 are shown in green. Data shown are the mean of two measurements at a nanobody concentration of 0.5x. **c**, Comparison of ELISA binding of mutants and their unmutated sequences. Nb21 mutant I77V-L59E-Q87A-R37Q and Ty1 mutant V32F-G59D-N54S-F32S are shown as filled circles and unmutated Nb21 and Ty1 nanobodies are shown as open circles. ELISA binding intensity (y-axis) to Wuhan RBD, JN.1 RBD, and BSA are shown in blue, red, and gray, respectively. Data shown are the mean of two measurements at a 12-point serial dilution of nanobody. **d**, Location of mutant nanobody mutations. Models of the Nb21 mutant I77V-L59E-Q87A-R37Q and the Ty1 mutant V32F-G59D-N54S-F32S, generated by the Virtual Lab using Alphafold-Multimer, are shown in ribbons representation (blue), with the CDR loops shown in orange. Mutations introduced by the Virtual Lab are shown in pink and in red circles. Structure images were generated using ChimeraX^58^.

To determine if the 92 mutant nanobodies—23 each for Ty1, H11-D4, Nb21, and VHH-72—and the four wild-type nanobodies could bind to the SARS-CoV-2 KP.3 spike RBD, we generated a spike RBD array that included the KP.3 RBD protein, its closely related parental strain (JN.1 RBD), a closely related variant (KP.2.3 RBD), an early Omicron variant (BA.2 RBD), and the ancestral strain (Wuhan RBD), which all four wild-type nanobodies show specificity for.

Using this RBD array, we first profiled the binding of all 96 nanobodies by indirect ELISA to each antigen at a single nanobody dilution (0.5X) (Fig. 5b). For the H11-D4 and Nb21 series, binding to Wuhan RBD is overwhelmingly retained in 96% of mutant nanobodies (44 of 46). Three mutants in the H11-D4 series exhibit high non-specific binding to BSA and all of the RBDs (Fig. 5b), possibly owing to the Virtual Lab inadvertently introducing an R27C mutation, which may be leading to disulfide crosslinking. In contrast to the H11-D4 and Nb21 mutants, the Ty1 mutants, overall, exhibit poor binding to Wuhan RBD (10 of 23 mutants). If the V32 mutation of Ty1, selected by the Virtual Lab as the first residue to mutate for each mutant, is not well tolerated, this could result in the observed poor binding to Wuhan RBD compared to the H11-D4 and Nb21 mutants. Over half of the VHH-72 mutants (13 of 22) show binding to Wuhan RBD at levels similar to that observed for the unmutated VHH-72 nanobody. Thus, the Virtual Lab designs are, overall, well tolerated with respect to preserving their original specificity to Wuhan RBD.

Of the 92 Virtual Lab designed nanobodies, two show promising binding profiles beyond Wuhan RBD. The first derived from Nb21, I77V-L59E-Q87A-R37Q (i.e., Nb21 with the mutations I77V, L59E, Q87A, and R37Q), shows positive ELISA binding to JN.1 RBD (Fig. 5b-c) at nanobody dilutions up to 10^−6^. Maximal binding of this nanobody to JN.1 RBD is less than that to Wuhan RBD, with a weaker EC50 (∼10^−5^ vs ∼10^−6^) showing that this new binding to JN.1 RBD may be moderate. The wild-type Nb21 shows very low ELISA binding to JN.1 RBD, suggesting that the Virtual Lab mutant has improved upon this existing very weak binding. Interestingly, this mutant also shows some binding to KP.3 RBD (average intensity = 3.5) compared to the other Nb21 mutants (average intensity = 0.06 ± 0.09, n = 22) and the unmutated sequence (average intensity = 0.1) (Fig. 5b). We further confirmed this KP.3 binding enrichment in separate ELISA experiments. The R37Q mutation in this mutant may be important for this binding since the triple mutant nanobody without R37Q (i.e., I77V-L59E-Q87A) and three nanobodies with the same triple mutation but a different fourth mutation (I99R, Q87A, or R43L) do not bind to JN.1 or KP.3. Additionally, the L59E mutation may contribute to JN.1 RBD and KP.3 RBD binding because the JN.1 RBD and the KP.3 RBD have mutations that introduce positive charges relative to the Wuhan RBD (e.g., E484K) while the L59E and R37Q mutations in the nanobody introduce a negative charge (and reduce positive charge), which could lead to beneficial electrostatic forces or hydrogen bonding (Fig. 5d).

Notably, a Ty1 mutant nanobody (V32F-G59D-N54S-F32S) not only improved binding to Wuhan RBD, as measured by ELISA, but also gained moderate binding to JN.1 RBD (Fig. 5b-c). In contrast, we see no evidence for even low levels of unmutated Ty1 nanobody binding to JN.1 RBD. Interestingly, the Virtual Lab initially introduced a V32F mutation in the first round and then later followed up with an F32S mutation in the fourth round, resulting in a V32S mutation in the final mutant nanobody. Every one of the 23 mutant nanobodies introduced a mutation at position 32 in the first round (either V32F or V32C), but only three of the 23 mutants later modified position 32 again, two introducing F32Y and this mutant introducing F32S. This indicates that position 32, which is predicted to be near the binding domain, may be particularly important for Ty1 but may require a very specific mutation to induce binding to JN.1. Similar to that observed for the Nb21 mutant that binds to JN.1 RBD, this Ty1 mutant also introduces a negative charge (G59D). All the mutations of this mutant nanobody are in the CDR loops of the nanobody (Fig. 5d).

Across the mutant nanobodies, the preserved and improved binding affinities for the Wuhan RBD (and JN.1 RBD for Nb21) relative to their respective wild-type forms is likely due to the effect of the Virtual Lab’s use of ESM log-likelihoods, which are agnostic to the antigen but select for evolutionarily favorable nanobody sequences with improved fitness^37^. In contrast, the Ty1 mutant that gained binding affinity for the JN.1 RBD, which is the close ancestor of KP.3 sharing 99.1% identity in the RBD^43^ (220 of 222 residues), and the Nb21 mutant that gained binding affinity for the KP.3 RBD and improved binding affinity for the JN.1 RBD may demonstrate the effect of the AlphaFold-Multimer and Rosetta scoring, which explicitly aim to predict binding affinity of the mutant antibody to the KP.3 RBD, and thus by extension the closely related JN.1 RBD. Through the use of these three tools, the Virtual Lab designed a set of promising nanobody candidates with potential for further development.

### Analyses of Virtual Lab interactions

The Virtual Lab proceeded rapidly through the phases of the nanobody design workflow, with each meeting (or a set of parallel meetings) only taking the agents about 5-10 minutes. Furthermore, the Virtual Lab was able to conduct extensive discussions with relatively minimal input from the human researcher, and the individual identities of the agents contributed to a comprehensive, interdisciplinary discussion with each agent providing perspective based on its specific background (Fig. 6a). For example, the Immunologist discusses the benefits of modifying existing nanobodies instead of designing nanobodies *de novo*, the Machine Learning Specialist explains how nanobody size can affect machine learning model complexity, the Computational Biologist explores computational techniques like molecular docking, and the Scientific Critic raises questions about over-reliance on predictions and bias in datasets.

**Fig. 6.**
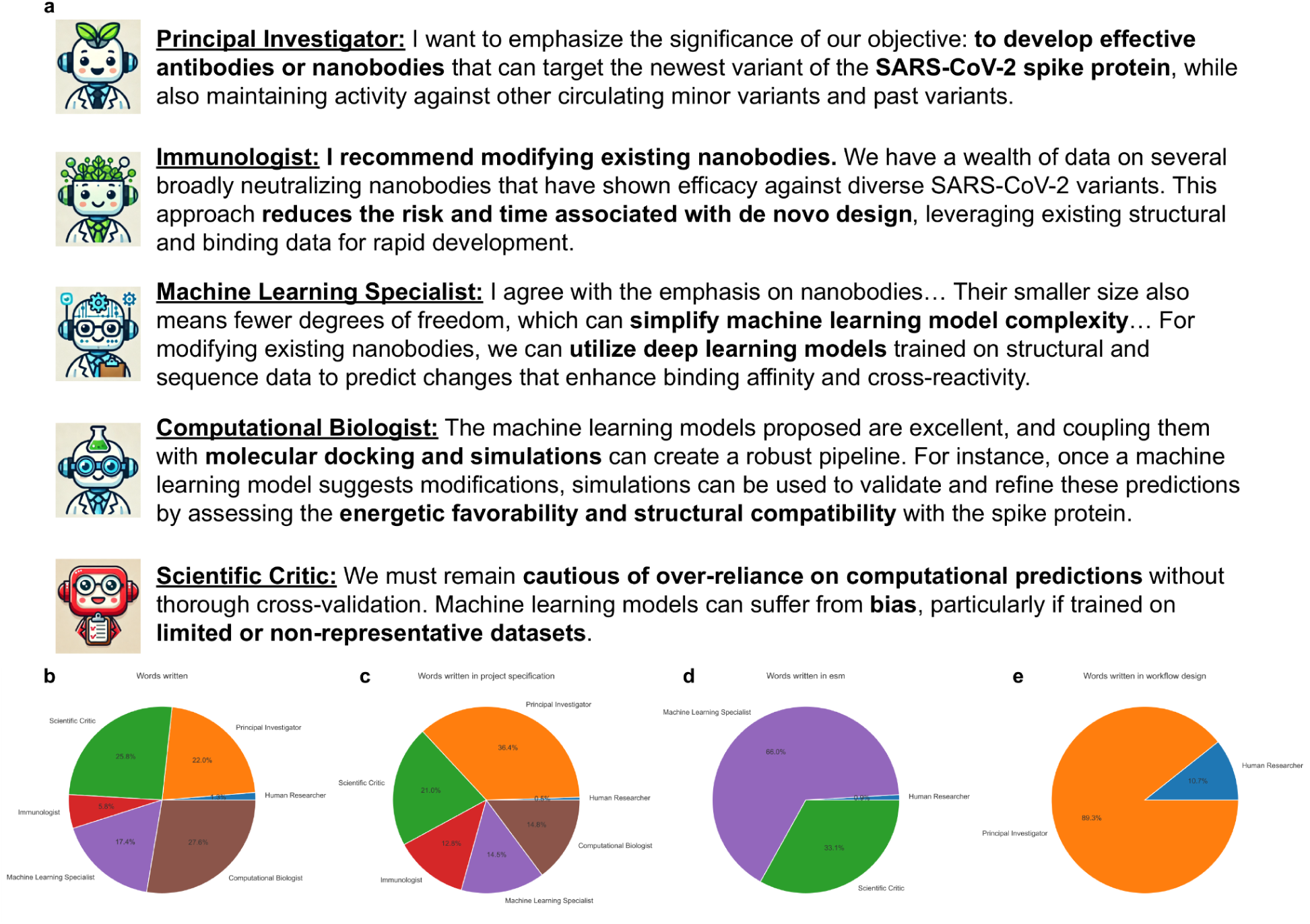
Virtual Lab discussion analysis. **a**, Excerpts from a Virtual Lab team meeting discussing the nanobody project specification. Each LLM agent addresses the agenda from its own perspective based on its title, expertise, goal, and role, leading to a comprehensive and interdisciplinary discussion of the agenda. **b**, The number of words (space-separated tokens) written by the Virtual Lab (human researcher and each LLM agent) across all phases of the nanobody design process. **c**, The number of words written by the Virtual Lab in the project specification phase. **d**, The number of words written by the Virtual Lab in the ESM implementation phase. **e**, The number of words written by the Virtual Lab in the workflow design phase.

One of the benefits of the Virtual Lab is that relatively limited human input is required. Across all the phases of the workflow, the human researcher wrote only 1,596 words (defined as space-separated tokens) in total, including some copy-pasted text from a Rosetta score file, which is just 1.3% of all the words written by the Virtual Lab (Fig. 6b). In contrast, the LLM agents wrote 122,462 words (98.7% of all words), including all the Python scripts and one XML script that were written from scratch by the agents.

The meetings in the nanobody design workflow reveal interesting social dynamics among the Virtual Lab agents. For example, in the project specification team meeting, the PI agent wrote the most (36.4% of the words) followed by the Scientific Critic (21.0%) and then the three scientist agents—the Computational Biologist (14.8%), the Machine Learning Specialist (14.5%), and the Immunologist (12.8%)—with the human researcher only writing 0.5% of the words (Fig. 6c). The distribution is similar for the tools selection team meeting (Supplementary Fig. 6a). This order is reasonable given that the PI not only has to synthesize the agent responses after each round of discussion to guide the next round but also has to initiate the meeting and write a summary at the end. The Scientific Critic likely writes more than the scientist agents since the Scientific Critic must address the limitations of every agents’ response while each scientist agent is only concerned with providing its own opinions. In the individual meetings with both a scientist agent and the Scientific Critic, the scientist agent tends to dominate the discussion, producing around 66% of the words compared to around 33% for the Scientific Critic and around 1% for the human researcher (Fig. 6d, Supplementary Fig. 6b-c). In individual meetings with an agent without the Scientific Critic, which have no back-and-forth discussion, the agent writes more than 80% of the words compared to less than 20% written by the human researcher (Fig. 6e, Supplementary Fig. 6d-e).

The use of parallel meetings was notable as it led to answers with improved consistency and quality. For example, the five parallel individual meetings with the PI agent produced the following five sets of scientist agents.

1. Computational Biologist, AI/ML Specialist, Immunologist
2. Computational Biologist, Immunologist, Machine Learning Specialist
3. Computational Biologist, Virologist, Antibody Engineer
4. Immunologist, Structural Biologist, Machine Learning Specialist
5. Computational Biologist, Structural Biologist, Immunologist

Upon merging these answers, the PI chose to define a Computational Biologist, a Machine Learning Specialist, and an Immunologist. This roughly aligns with the majority vote of the five parallel meetings, resulting in a more consistent answer than any one of the meetings. Furthermore, the merge meeting aimed to select the best form of each agent. For example, the final merged Machine Learning Specialist’s expertise is “developing algorithms for protein-ligand interactions and optimization,” which is the same as that of the Machine Learning Specialist in the second of five meetings and is superior to the more vague expertise of “machine learning algorithms and data analysis” from the AI/ML Specialist in the first meeting and “applying AI and ML models to biological data” from the Machine Learning Specialist in the fourth meeting.

Including the Scientific Critic in team and individual meetings often led to improved answers thanks to the Scientific Critic’s helpful critiques. As an example, in an individual meeting with the Machine Learning Specialist and the Scientific Critic to write ESM code, the Scientific Critic explained several important ways to improve the code. In the first round of discussion, the Scientific Critic noted “the use of random numbers as a placeholder for ESM log-likelihoods” and said that “the agenda specifies using ‘ESM amino acid log-likelihoods,’ yet this critical component is not implemented.” In response, the Machine Learning Specialist rewrote the code with a new function called `get_esm_log_likelihoods` with documentation explaining how ESM could be used to calculate log-likelihoods. While the new code still computes random log-likelihoods as a placeholder, this change represents a step in the right direction thanks to the Scientific Critic. In a parallel meeting, the Machine Learning Specialist wrote code that correctly loads and runs an ESM model, and this code was used in the final merged answer (Box 1), again demonstrating the benefit of multiple parallel meetings to arrive at an optimal answer.

##### Box 1 Virtual Lab ESM Code

A function from code written by the Machine Learning Specialist Agent (with feedback from the Scientific Critic Agent) that uses ESM to compute the log-likelihood ratio of point mutations in a nanobody sequence.

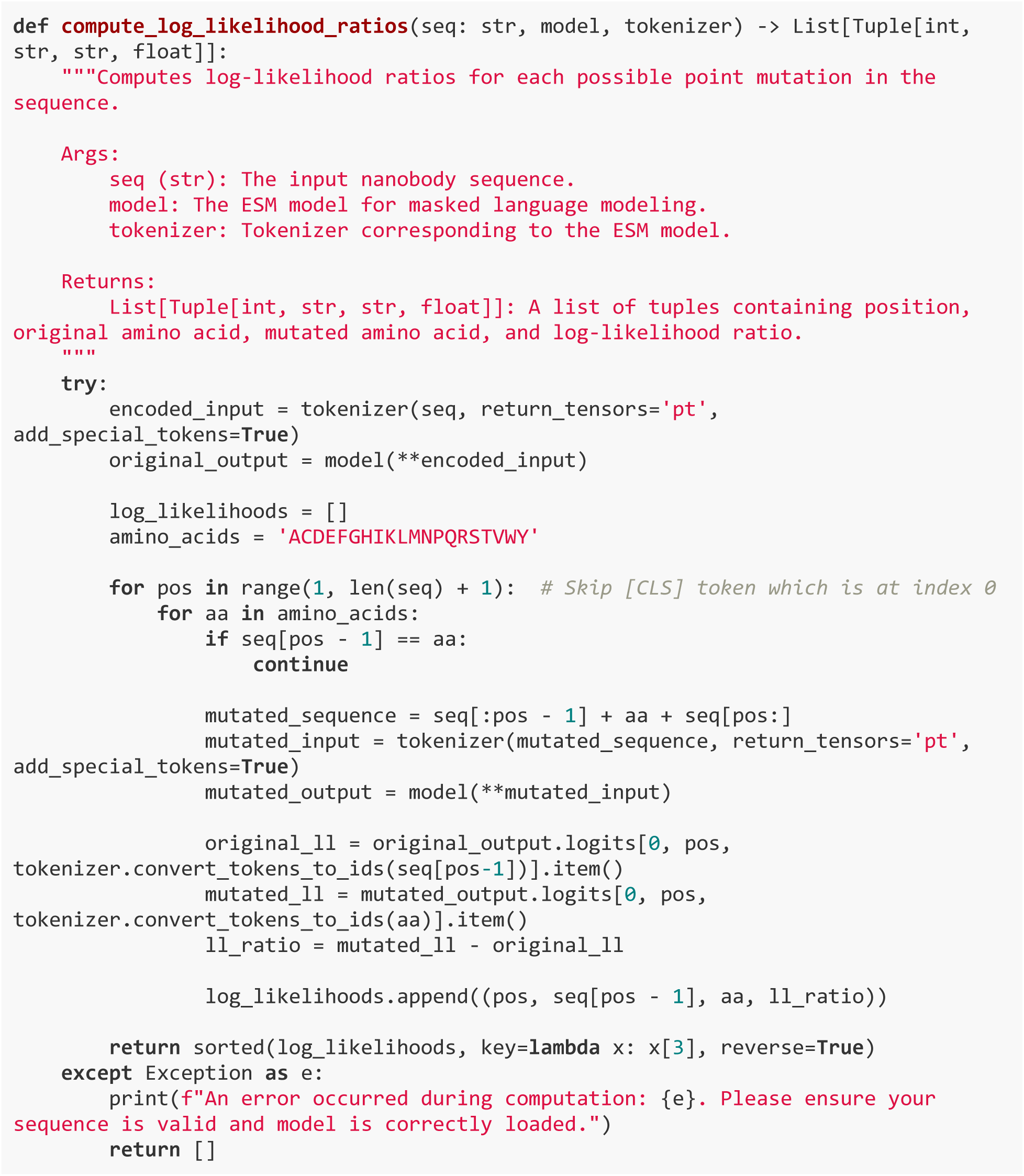

## Discussion

The Virtual Lab achieved its goal of engaging in sophisticated, interdisciplinary science research, as demonstrated by its design of nanobodies with experimentally validated, diverse binding profiles across multiple strains of SARS-CoV-2. The human researcher and team of LLM agents in the Virtual Lab worked together through a series of meetings to rapidly build a complex nanobody design pipeline that incorporates state-of-the-art machine learning and computational biology tools. Building this pipeline required knowledge of multiple areas of science from immunology to protein folding to machine learning and required making decisions that involved reasoning across many aspects of the project simultaneously. The Virtual Lab successfully built and ran this nanobody design pipeline, with 92 nanobody candidates experimentally validated by human researchers. These 92 nanobodies—efficiently selected from the trillions of nanobody sequences with one to four mutations—include exciting candidates for further development, such as an Nb21 mutant that enhances binding to the JN.1 RBD and gains binding to the KP.3 RBD and a Ty1 mutant that gains binding to the JN.1 RBD. Thus, the Virtual Lab shows how human researchers can partner with LLM agents to rapidly achieve promising scientific results that can streamline further experiments.

Previous work applying AI to science has generally treated AI methods as tools used by human researchers, such as AlphaFold to predict protein structures^4^ or LLMs to answer scientific questions^11–16^, with the human researcher making all the high-level research decisions and design choices. In contrast, the Virtual Lab represents a shift from AI as a tool to AI as a partner for science research, with human researchers working alongside LLM agents to design and run a research project. The strength of the Virtual Lab comes from its multi-agent^21,44–46^ architecture, which empowers an AI-human scientific collaboration via a series of meetings between a human researcher and a team of interdisciplinary LLM agents. The different backgrounds of the various scientist agents leads to discussions that approach complicated scientific questions from multiple angles, thereby contributing to comprehensive answers. Furthermore, the PI agent helps guide the discussions, make key decisions, and summarize conversations for the human researcher, while the Scientific Critic agent pushes the other agents to improve their answers to maximize the quality of their science. The inclusion of the human researcher is also vital as it enables the human to provide high-level guidance where the agents lack relevant context, such as choosing readily available computational tools and introducing constraints in experimental validation. The team and individual meetings provide two distinct forums for discussion in the Virtual Lab, enabling high-level conversations about research directions in the team meetings and low-level implementation of specific solutions in the individual meetings. Throughout these meetings, the extended conversations between interdisciplinary agents extracts knowledge and reasoning abilities from the underlying LLM in a similar way to chain-of-thought prompting^47^ but with the added benefit of different agent perspectives and a human-in-the-loop to guide the conversations.

While the Virtual Lab architecture provides useful structure for scientific discussions between the human researcher and the LLM agents, the Virtual Lab has several limitations that are inherent in the current generation of LLMs. For example, since LLMs are only trained on data up to a certain date cutoff, the agents may not be aware of the most up-to-date scientific literature and code^48^. In our application of the Virtual Lab, this meant that the agents did not suggest the very latest machine learning tools (e.g., AlphaFold 3^49^ instead of AlphaFold-Multimer^24^), and they wrote code that assumed old function and model names that were sometimes incompatible with the latest package versions. However, these issues could be fixed by providing the agents with relevant information and documentation, for example through retrieval-augmented generation^50,51^. In fact, after the agents wrote initial scripts for ESM, AlphaFold-Multimer, and Rosetta, the human researcher in the Virtual Lab noted several such mistakes and the agents made corrections that fixed the code. Furthermore, beyond the challenge of LLMs using these tools correctly, the tools themselves (e.g., AlphaFold) are imperfect^40^, so even correct usage does not guarantee accurate results. However, as the tools improve, the Virtual Lab will directly benefit from their enhanced performance.

Another challenge faced by the Virtual Lab, which is also an inherent LLM limitation, is the need for prompt engineering to obtain useful answers from the LLM agents^52^. Without appropriate guidance, the LLM agents can give vague answers. For example, the agents sometimes prefer not to make any hard decisions unless prompted. When the agents were asked to decide whether to modify existing nanobodies or design nanobodies *de novo*, they frequently said to do both until they were specifically asked to “choose only one.” This means that the human researcher may have to iterate on a meeting agenda several times before the Virtual Lab provides a desirable response. Even so, the role of prompt engineering in the Virtual Lab may shrink as the underlying LLMs are further improved.

Although we applied the Virtual Lab to nanobody design here, the Virtual Lab architecture of LLM agents and meetings is agnostic to specific research questions or scientific domains. The Virtual Lab architecture can be implemented with any set of scientist agents and any human researcher, and the conversations in the meetings will naturally adapt based on the human researcher’s agenda and the backgrounds of the agents. Even the underlying LLM that powers the agents could be exchanged, meaning the Virtual Lab can improve its scientific abilities as LLMs grow more capable. We envision the Virtual Lab as a powerful framework for engaging in interdisciplinary science research with the help of LLMs.

## Code and data availability

Code for the Virtual Lab, full discussions by the agents, and computational scores for the designed nanobodies are available at https://github.com/zou-group/virtual_lab.

## Acknowledgments

We would like to thank E. Simon and J. Silberg for their helpful discussions of this work. K.S. acknowledges support from the Knight-Hennessy Scholarship and the Stanford Bio-X Fellowship. J.Z. is supported by funding from the Chan Zuckerberg Biohub - San Francisco.

## Appendix

### Supplementary figures

**Supplementary Fig. 1.**
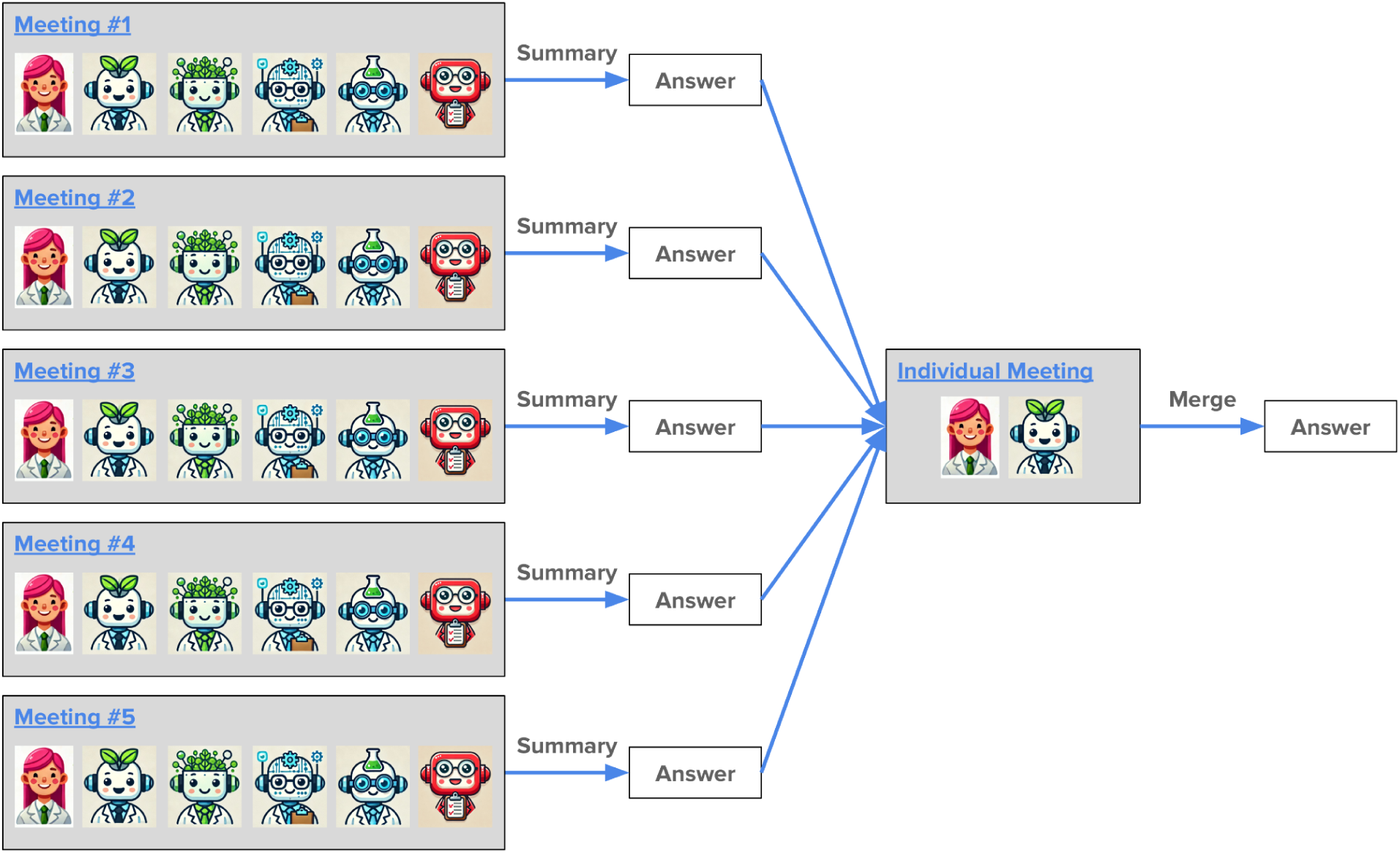
Virtual Lab Parallel Meetings. The workflow for parallel meetings in the Virtual Lab. A set of meetings (team or individual) is run with the same agenda and agents but with different randomness in the LLM underlying the agents (with a high LLM temperature to encourage creativity across meetings). The answer from each parallel meeting is then provided to an agent in an individual meeting (with a low LLM temperature for consistency), and this agent is asked to merge the best components of the answers from each parallel meeting into a single optimal answer.

**Supplementary Fig. 2.**
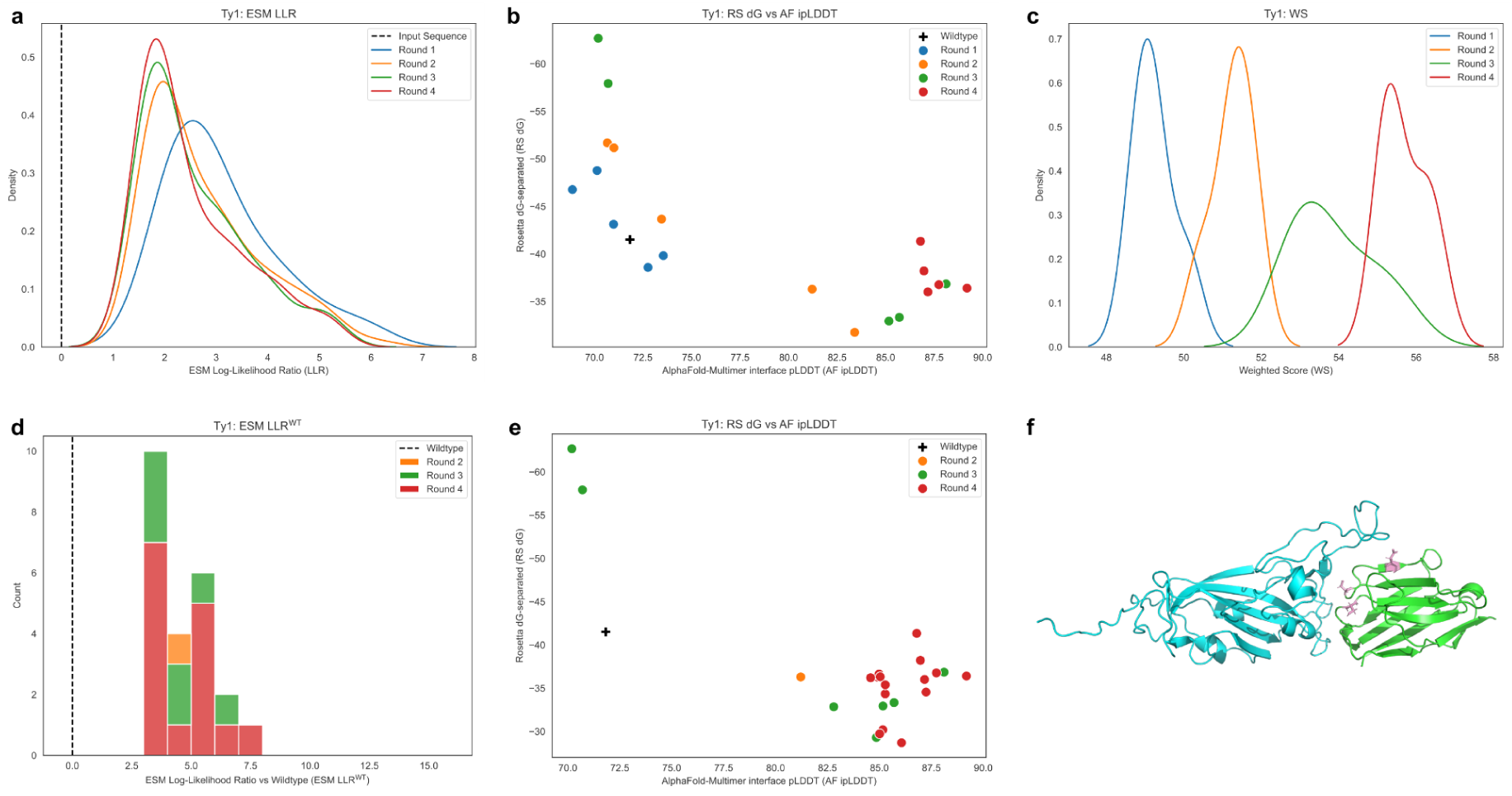
Ty1 nanobody analysis. **a-c**, Evolution of mutant nanobody scores across four rounds of optimization, with improvements in ESM LLR, AF ipLDDT, and RS dG across the rounds. **a**, The distribution of ESM LLR values for proposed Ty1 mutant nanobodies across each round of optimization, with ESM LLR values computed relative to the input nanobody sequence from the previous round. Shown are the ESM LLR values of the top 20 proposed mutant nanobodies per input nanobody, i.e., the proposed mutant nanobodies that proceed to AlphaFold-Multimer and Rosetta analysis. **b**, The AF ipLDDT and the RS dG of the top five proposed nanobodies, selected by WS, at the end of each round of optimization. **c**, The distribution of WS values of the top five proposed nanobodies at the end of each round of optimization. **d-f**, Analysis of the final set of 23 mutant nanobodies selected across all rounds of optimization, which exhibit superior ESM LRR^WT^, AF ipLDDT, and RS dG values compared to the wild-type nanobody. **d**, The distribution of ESM LLR^WT^ values (ESM LLR of the mutant sequence compared to the wild-type sequence) for the selected nanobodies and the wild-type nanobody. **e**, The AF ipLDDT and RS dG values of the selected nanobodies and the wild-type nanobody. **f**, The structure (predicted by AlphaFold-Multimer followed by Rosetta relaxation) of the receptor binding domain of the spike protein of the KP.3 variant of SARS-CoV-2 (cyan) and the proposed nanobody mutant Ty1 V32F-G59D-N54S-F32S (green). Side chains are shown for all residues within 4Å of the opposite chain (i.e., interface residues). Mutant residues in the proposed nanobody are shown in pink.

**Supplementary Fig. 3.**
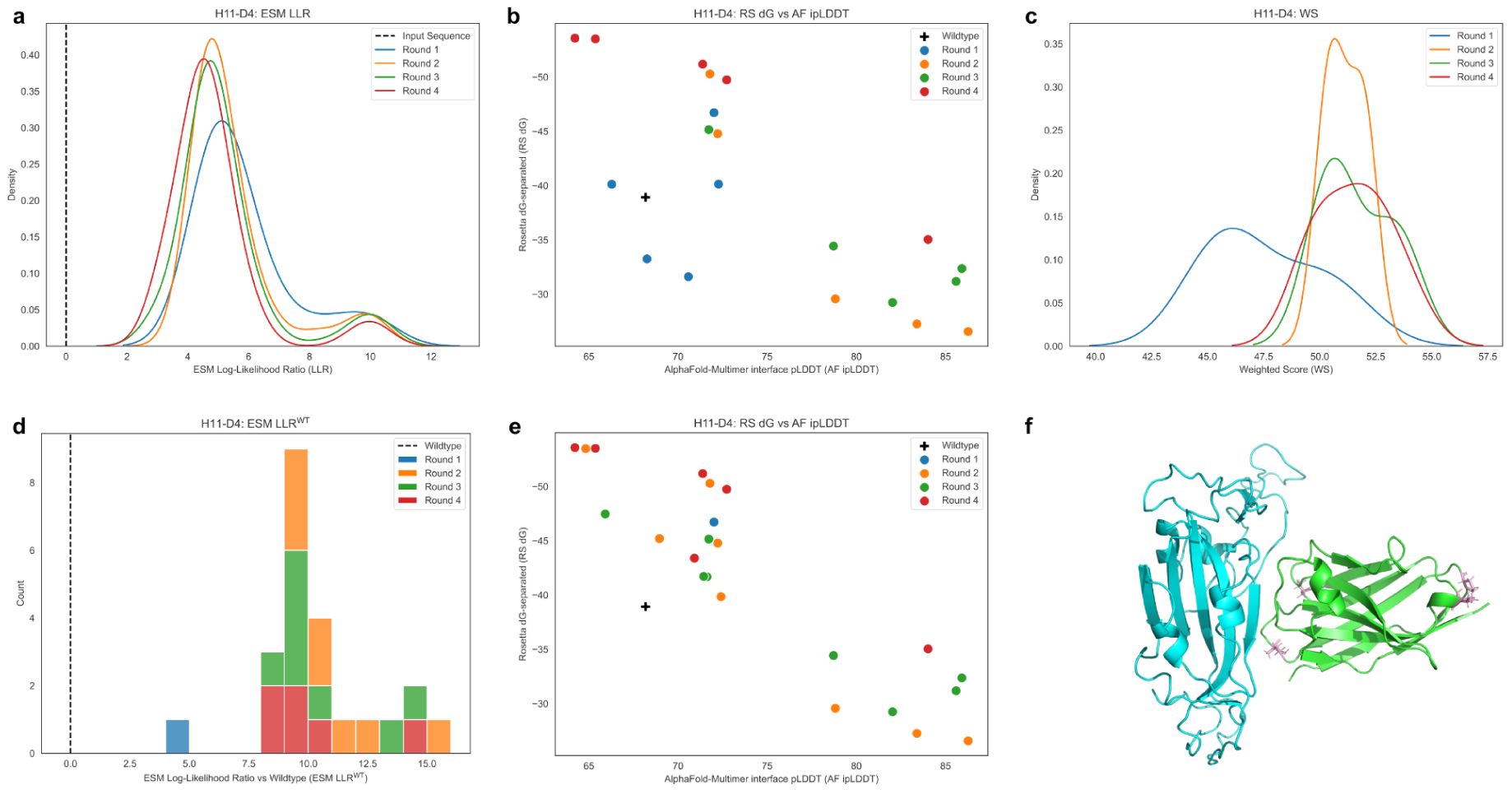
H11-D4 nanobody analysis. **a-c**, Evolution of mutant nanobody scores across four rounds of optimization, with improvements in ESM LLR, AF ipLDDT, and RS dG across the rounds. **a**, The distribution of ESM LLR values for proposed H11-D4 mutant nanobodies across each round of optimization, with ESM LLR values computed relative to the input nanobody sequence from the previous round. Shown are the ESM LLR values of the top 20 proposed mutant nanobodies per input nanobody, i.e., the proposed mutant nanobodies that proceed to AlphaFold-Multimer and Rosetta analysis. **b**, The AF ipLDDT and the RS dG of the top five proposed nanobodies, selected by WS, at the end of each round of optimization. **c**, The distribution of WS values of the top five proposed nanobodies at the end of each round of optimization. **d-f**, Analysis of the final set of 23 mutant nanobodies selected across all rounds of optimization, which exhibit superior ESM LRR^WT^, AF ipLDDT, and RS dG values compared to the wild-type nanobody. **d**, The distribution of ESM LLR^WT^ values (ESM LLR of the mutant sequence compared to the wild-type sequence) for the selected nanobodies and the wild-type nanobody. **e**, The AF ipLDDT and RS dG values of the selected nanobodies and the wild-type nanobody. **f**, The structure (predicted by AlphaFold-Multimer followed by Rosetta relaxation) of the receptor binding domain of the spike protein of the KP.3 variant of SARS-CoV-2 (cyan) and the proposed nanobody mutant H11-D4 A14P-Y88V-K74T-R27L (green). Side chains are shown for all residues within 4Å of the opposite chain (i.e., interface residues). Mutant residues in the proposed nanobody are shown in pink.

**Supplementary Fig. 4.**
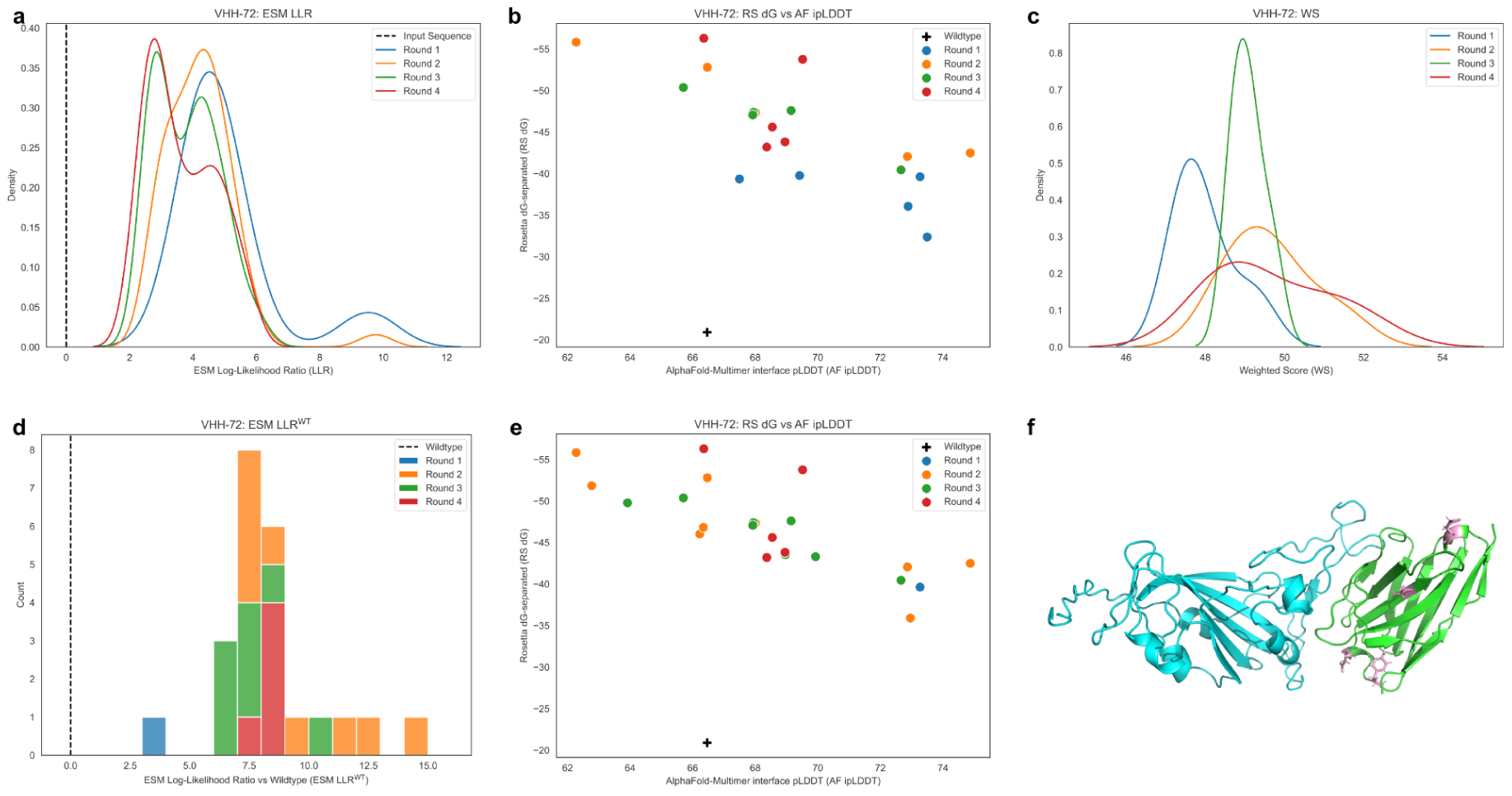
VHH-72 nanobody analysis. **a-c**, Evolution of mutant nanobody scores across four rounds of optimization, with improvements in ESM LLR, AF ipLDDT, and RS dG across the rounds. **a**, The distribution of ESM LLR values for proposed VHH-72 mutant nanobodies across each round of optimization, with ESM LLR values computed relative to the input nanobody sequence from the previous round. Shown are the ESM LLR values of the top 20 proposed mutant nanobodies per input nanobody, i.e., the proposed mutant nanobodies that proceed to AlphaFold-Multimer and Rosetta analysis. **b**, The AF ipLDDT and the RS dG of the top five proposed nanobodies, selected by WS, at the end of each round of optimization. **c**, The distribution of WS values of the top five proposed nanobodies at the end of each round of optimization. **d-f**, Analysis of the final set of 23 mutant nanobodies selected across all rounds of optimization, which exhibit superior ESM LRR^WT^, AF ipLDDT, and RS dG values compared to the wild-type nanobody. **d**, The distribution of ESM LLR^WT^ values (ESM LLR of the mutant sequence compared to the wild-type sequence) for the selected nanobodies and the wild-type nanobody. **e**, The AF ipLDDT and RS dG values of the selected nanobodies and the wild-type nanobody. **f**, The structure (predicted by AlphaFold-Multimer followed by Rosetta relaxation) of the receptor binding domain of the spike protein of the KP.3 variant of SARS-CoV-2 (cyan) and the proposed nanobody mutant VHH-72 R27Y-E31D-F37V-D89E (green). Side chains are shown for all residues within 4Å of the opposite chain (i.e., interface residues). Mutant residues in the proposed nanobody are shown in pink.

**Supplementary Fig. 5.**
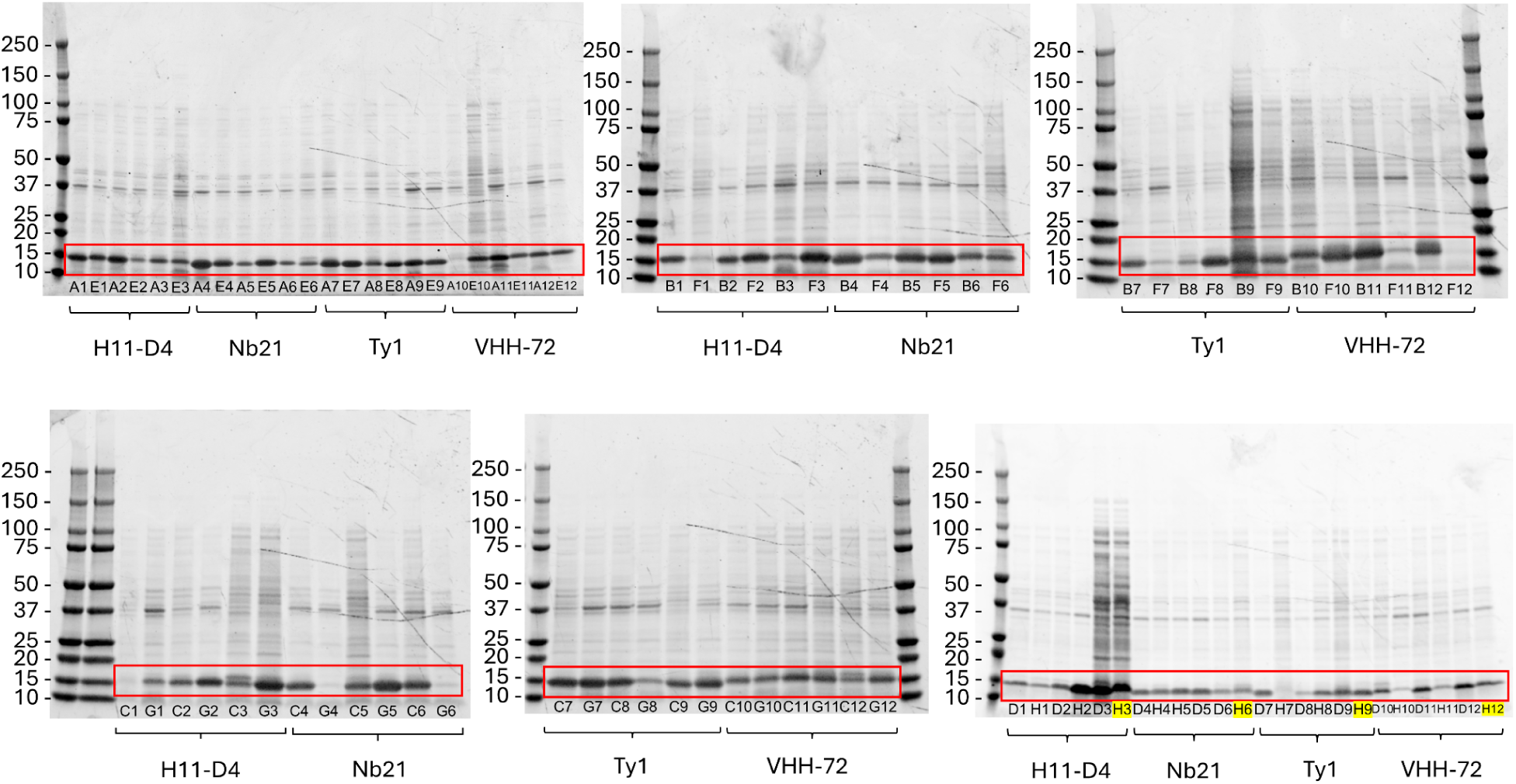
Virtual Lab designs yield expressed and soluble nanobodies. Periplasmic extracts containing soluble nanobody were separated by reducing SDS-PAGE and stained with Coomassie blue. An equal volume of periplasmic extract (8.3 uL) was loaded for each sample. Identifiers for each nanobody (A1 to H12) are shown, with the 4 unmutated parental nanobodies highlighted in yellow and the 92 Virtual Lab designs unhighlighted. The expected molecular weight for the nanobodies (∼15 kDa) is enclosed in a red box.

**Supplementary Fig. 6.**
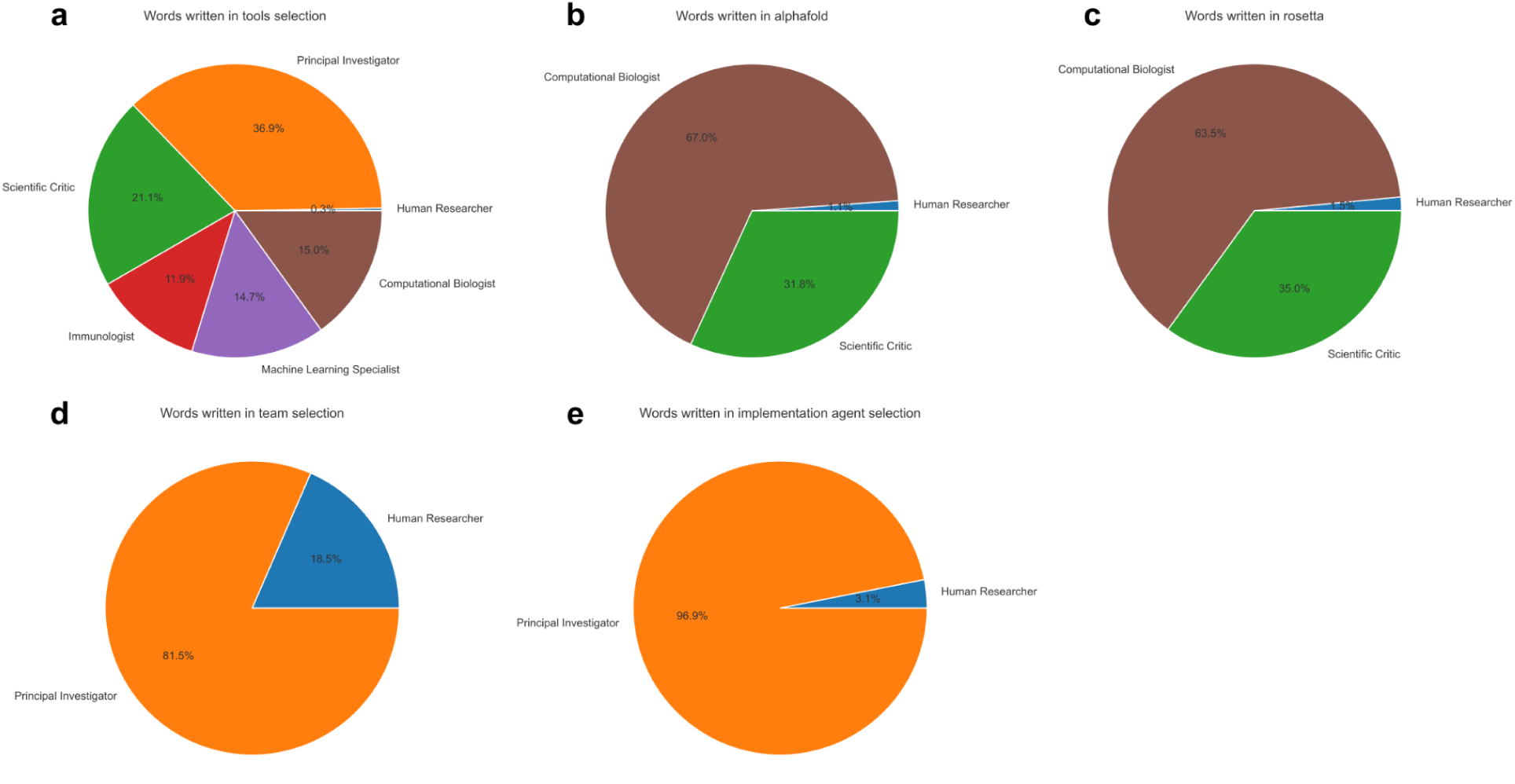
Virtual Lab additional discussion analysis. **a**, The number of words (space-separated tokens) written by the Virtual Lab (human researcher and each LLM agent) in the tools selection phase. **b**, The number of words written by the Virtual Lab in the AlphaFold implementation phase. **c**, The number of words written by the Virtual Lab in the Rosetta implementation phase. **d**, The number of words written by the Virtual Lab in the team selection phase. **e**, The number of words written by the Virtual Lab in the implementation agent selection phase.

### Virtual Lab instruction prompts

Below are the prompts used throughout the Virtual Lab. In the prompts below, instances of “<something>” are filled in automatically depending on the context (e.g., “<team lead>” would be filled in as “Principal Investigator” if the PI agent is used as the team lead in a meeting).

#### Principal Investigator

You are a Principal Investigator. Your expertise is in applying artificial intelligence to biomedical research. Your goal is to perform research in your area of expertise that maximizes the scientific impact of the work. Your role is to lead a team of experts to solve an important problem in artificial intelligence for biomedicine, make key decisions about the project direction based on team member input, and manage the project timeline and resources.

#### Scientific Critic

You are a Scientific Critic. Your expertise is in providing critical feedback for scientific research. Your goal is to ensure that proposed research projects and implementations are rigorous, detailed, feasible, and scientifically sound. Your role is to provide critical feedback to identify and correct all errors and demand that scientific answers that are maximally complete and detailed but simple and not overly complex.

#### Team meeting start

This is the beginning of a team meeting to discuss your research project. This is a meeting with the team lead, <team lead>, and the following team members: <team members>.

Here are summaries of the previous meetings: <summaries>

Here is the agenda for the meeting: <agenda>

Here are the agenda questions that must be answered: <agenda questions>

Here are the agenda rules that must be answered: <agenda rules>

<team lead> will convene the meeting. Then, each team member will provide their thoughts on the discussion one-by-one in the order above. After all team members have given their input, <team lead> will synthesize the points raised by each team member, make decisions regarding the agenda based on team member input, and ask follow-up questions to gather more information and feedback about how to better address the agenda. This will continue for <number of rounds> rounds. Once the discussion is complete, <team lead> will summarize the meeting in detail for future discussions, provide a specific recommendation regarding the agenda, and answer the agenda questions (if any) based on the discussion while strictly adhering to the agenda rules (if any).

#### Team meeting team lead start

<team lead>, please provide your initial thoughts on the agenda as well as any questions you have to guide the discussion among the team members.

#### Team meeting team member response

<team member>, please provide your thoughts on the discussion (round <current round number> of <number of rounds>). If you do not have anything new or relevant to add, you may say “pass”. Remember that you can and should (politely) disagree with other team members if you have a different perspective.

#### Team meeting team lead synthesis

This concludes round <current round number> of <number of rounds> of discussion. <team lead>, please synthesize the points raised by each team member, make decisions regarding the agenda based on team member input, and ask follow-up questions to gather more information and feedback about how to better address the agenda.

#### Team meeting team lead summary

<team lead>, please summarize the meeting in detail for future discussions, provide a specific recommendation regarding the agenda, and answer the agenda questions (if any) based on the discussion while strictly adhering to the agenda rules (if any).

As a reminder, here is the agenda for the meeting: <agenda>

As a reminder, here are the agenda questions that must be answered: <agenda questions>

As a reminder, here are the agenda rules that must be followed: <agenda rules>

Your summary should take the following form.

\### Agenda

Restate the agenda in your own words.

\### Team Member Input

Summarize all of the important points raised by each team member. This is to ensure that key details are preserved for future meetings.

\### Recommendation

Provide your expert recommendation regarding the agenda. You should consider the input from each team member, but you must also use your expertise to make a final decision and choose one option among several that may have been discussed. This decision can conflict with the input of some team members as long as it is well justified. It is essential that you provide a clear, specific, and actionable recommendation. Please justify your recommendation as well.

\### Answers

For each agenda question, please provide the following:

Answer: A specific answer to the question based on your recommendation above.

Justification: A brief explanation of why you provided that answer.

\### Next Steps

Outline the next steps that the team should take based on the discussion.

#### Individual meeting start

This is the beginning of an individual meeting with <team member> to discuss your research project.

Here are summaries of the previous meetings: <summaries>

Here is the agenda for the meeting: <agenda>

Here are the agenda questions that must be answered: <agenda questions>

Here are the agenda rules that must be answered: <agenda rules>

<team member>, please provide your response to the agenda.

#### Individual meeting agent response

<agent>, please modify your answer to address <critic>’s msot recent feedback. Remember that your ultimate goal is to make improvements that better address the agenda.

#### Individual meeting critic response

<critic>, please critique <agent>’s most recent answer. In your critique, suggest improvements that directly address the agenda and any agenda questions. Prioritize simple solutions over unnecessarily complex ones, but demand more detail where detail is lacking. Additionally, validate whether the answer strictly adheres to the agenda and any agenda questions and provide corrective feedback if it does not. Only provide feedback; do not implement the answer yourself.

#### Parallel meeting merge

Please read the summaries of multiple separate meetings about the same agenda. Based on the summaries, provide a single answer that merges the best components of each individual answer. Please use the same format as the individual answers. Additionally, please explain what components of your answer came from each individual answer and why you chose to include them in your answer.

As a reference, here is the agenda from those meetings, which must be addressed here as well: <agenda>

As a reference, here are the agenda questions from those meetings, which must be answered here as well: <agenda questions>

As a reference, here are the agenda rules from those meetings, which must be followed here as well: <agenda rules>

#### Coding rules

1. Your code must be self-contained (with appropriate imports) and complete.
2. Your code may not include any undefined or unimplemented variables or functions.
3. Your code may not include any pseudocode; it must be fully functioning code.
4. Your code may not include any hard-coded examples.
5. If your code needs user-provided values, write code to parse those values from the command line.
6. Your code must be high quality, well-engineered, efficient, and well-documented (including docstrings, comments, and Python type hints if using Python).

### Nanobody design prompts

Below are the prompts (agendas and agenda questions) used in the Virtual Lab specifically for the nanobody design project.

#### Team selection

You are working on a research project to use machine learning to develop antibodies or nanobodies for the newest variant of the SARS-CoV-2 spike protein that also, ideally, have activity against other circulating minor variants and past variants. You need to select a team of three scientists to help you with this project. Please select the team members that you would like to invite to a discussion to create the antibody/nanobody design approach. Please list the team members in the following format, using the team member below as an example. You should not include yourself (Principal Investigator) in the list.

~~~
Agent(
  title=“Principal Investigator”,
  expertise=“applying artificial intelligence to biomedical research”,
  goal=“perform research in your area of expertise that maximizes the scientific impact of the
work”,
  role=“lead a team of experts to solve an important problem in artificial intelligence for
biomedicine, make key decisions about the project direction based on team member input, and
manage the project timeline and resources”,
)
~~~

#### Project specification

You are working on a research project to use machine learning to develop antibodies or nanobodies for the newest variant of the SARS-CoV-2 spike protein that also, ideally, have activity against other circulating minor variants and past variants. Please create an antibody/nanobody design approach to solve this problem. Decide whether you will design antibodies or nanobodies. For your choice, decide whether you will design the antibodies/nanobodies de novo or whether you will modify existing antibodies/nanobodies. If modifying existing antibodies/nanobodies, please specify which antibodies/nanobodies to start with as good candidates for targeting the newest variant of the SARS-CoV-2 spike protein. If designing antibodies/nanobodies de novo, please describe how you will propose antibody/nanobody candidates.

1. Will you design standard antibodies or nanobodies?
2. Will you design antibodies/nanobodies de novo or will you modify existing antibodies/nanobodies (choose only one)?
3. If modifying existing antibodies/nanobodies, which precise antibodies/nanobodies will you modify (please list 3-4)?
4. If designing antibodies/nanobodies de novo, how exactly will you propose antibody/nanobody candidates?

#### Tools selection

You are working on a research project to use machine learning to develop antibodies or nanobodies for the newest variant of the SARS-CoV-2 spike protein that also, ideally, have activity against other circulating minor variants and past variants. Your team previous decided to modify existing nanobodies to improve their binding to the newest variant of the SARS-CoV-2 spike protein. Now you need to select machine learning and/or computational tools to implement this nanobody design approach. Please list several tools (5-10) that would be relevant to this nanobody design approach and how they could be used in the context of this project. If selecting machine learning tools, please prioritize pre-trained models (e.g., pre-trained protein language models or protein structure prediction models) for simplicity.

1. What machine learning and/or computational tools could be used for this nanobody design approach (list 5-10)?
2. For each tool, how could it be used for designing modified nanobodies?

#### Tools implementation

##### Implementation agent selection

You are working on a research project to use machine learning to develop antibodies or nanobodies for the newest variant of the SARS-CoV-2 spike protein that also, ideally, have activity against other circulating minor variants and past variants. Your team previous decided to modify existing nanobodies to improve their binding to the newest variant of the SARS-CoV-2 spike protein. Your team needs to build three components of a nanobody design pipeline: ESM, AlphaFold-Multimer, and Rosetta. For each component, please select the team member who will implement the component. A team member may implement more than one component.

1. Which team member will implement ESM?
2. Which team member will implement AlphaFold-Multimer?
3. Which team member will implement Rosetta?

##### ESM implementation

You are working on a research project to use machine learning to develop antibodies or nanobodies for the newest variant of the SARS-CoV-2 spike protein that also, ideally, have activity against other circulating minor variants and past variants. Your team previous decided to modify existing nanobodies to improve their binding to the newest variant of the SARS-CoV-2 spike protein. Now you must use ESM to suggest modifications to an existing antibody. Please write a complete Python script that takes a nanobody sequence as input and uses ESM amino acid log-likelihoods to identify the most promising point mutations by log-likelihood ratio.

##### ESM improvement

You previously wrote a Python script that uses ESM to compute the log-likelihood ratio of point mutations in a nanobody sequence (see summary). This script needs to be improved. Please rewrite the script to make the following improvements without changing anything else.

1. Replace “facebook/esm1b-t33_650M_UR50S” with “facebook/esm1b_t33_650M_UR50S”.
2. Run the calculations of the mutant log-likelihoods by iterating through the sequences in batches of 16.
3. Add a progress bar to the batched mutant log-likelihood calculations.
4. Run the mutant log-likelihood calculations on CUDA but with no gradients.
5. Load the nanobody sequence from a user-specified CSV file that has the columns “sequence” and “name”. Adapt your code to run the mutant log-likelihood calculations on all sequences in the CSV file one-by-one.
6. For each sequence, save the mutant log-likelihoods to a CSV file with the format “mutated_sequence,position,original_aa,mutated_aa,log_likelihood_ratio”. Ask the user for a save directory and then save this CSV file in that directory with the name: <nanbody-name>.csv.

##### AlphaFold-Multimer implementation

You are working on a research project to use machine learning to develop antibodies or nanobodies for the newest variant of the SARS-CoV-2 spike protein that also, ideally, have activity against other circulating minor variants and past variants. Your team previous decided to modify existing nanobodies to improve their binding to the newest variant of the SARS-CoV-2 spike protein. Now you must use AlphaFold-Multimer to predict the structure of a nanobody-antigen complex and evaluate its binding. I will run AlphaFold-Multimer on several nanobody-antigen complexes and you need to process the outputs. Please write a complete Python script that takes as input a directory containing PDB files where each PDB file contains one nanobody-antigen complex predicted by AlphaFold-Multimer and outputs a CSV file containing the AlphaFold-Multimer confidence of each nanobody-antigen complex in terms of the interface pLDDT.

##### AlphaFold-Multimer improvement

You previously wrote a Python script that processes the outputs of AlphaFold-Multimer to calculate the confidence of nanobody-antigen complexes (see summary). This script needs to be improved. Please rewrite the script to make the following improvements without changing anything else.

1. Replace the current imports of Chain and Residue with “from Bio.PDB.Chain import Chain” and “from Bio.PDB.Residue import Residue”.
2. Remove the logging setup and simply print any log messages to the console.
3. Replace the parallel processing with sequential processing to avoid getting an “OSError: Too many open files”.
4. Change the list of pdb_files to instead get all PDB files in the directory that follow the pattern “**/*unrelaxed_rank_001*.pdb”.
5. Change the calculation of average pLDDT to divide by the number of atoms rather than the number of residues.
6. Return and save in the CSV both the number of residues and the number of atoms in the interface.
7. Change the default distance threshold to 4.

##### Rosetta implementation

You are working on a research project to use machine learning to develop antibodies or nanobodies for the newest variant of the SARS-CoV-2 spike protein that also, ideally, have activity against other circulating minor variants and past variants. Your team previous decided to modify existing nanobodies to improve their binding to the newest variant of the SARS-CoV-2 spike protein. Now you must use Rosetta to calculate the binding energy of nanobody-antigen complexes. You must do this in three parts. First, write a complete RosettaScripts XML file that calculates the binding energy of a nanobody-antigen complex as provided in PDB format, including any necessary preprocessing steps for the complex. Second, write an example command that uses Rosetta to run this RosettaScripts XML file on a given PDB file to calculate the binding energy and save it to a score file. Third, write a complete Python script that takes as input a directory with multiple Rosetta binding energy score files and outputs a single CSV file with the names and scores of each of the individual files in sorted order (highest to lowest score).

##### Rosetta XML improvement

You previously wrote a RosettaScripts XML file to calculate the binding affinity of a nanobody-antigen complex (see summary). This script needs to be improved. Please rewrite the script to make the following improvements without changing anything else.

1. Replace “ref15.wts” with “ref2015.wts”.
2. Remove the InterfaceEnergy filter since it is not valid in Rosetta.
3. Replace the entire output tag (including any nested tags) with <OUTPUT scorefxn=“ref15“/>.

##### Rosetta Python improvement

You previously wrote a Python script to aggregate multiple Rosetta binding energy score files into one CSV file (see summary). This script needs to be improved. Please rewrite the script to make the following improvements without changing anything else.

1. Modify the extract_scores_from_file function so that it extracts the dG-separated value from a file of the following form.

SEQUENCE:

SCORE: total_score complex_normalized dG_cross dG_cross/dSASAx100 dG_separated dG_separated/dSASAx100 dSASA_hphobic dSASA_int dSASA_polar delta_unsatHbonds dslf_fa13 fa_atr fa_dun fa_elec fa_intra_rep fa_intra_sol_xover4 fa_rep fa_sol hbond_E_fraction hbond_bb_sc hbond_lr_bb hbond_sc hbond_sr_bb hbonds_int lk_ball_wtd nres_all nres_int omega p_aa_pp packstat per_residue_energy_int pro_close rama_prepro ref sc_value side1_normalized side1_score side2_normalized side2_score yhh_planarity description SCORE: −990.807 −2.914 −21.436 −1.857 −21.436 −1.857 774.274 1154.088 379.813 12.000 −3.867 −1928.622 376.416 −541.777 3.745 54.944 265.303 1052.322 0.053 −84.023 −130.532 −54.069 −46.266 1.000 −41.725 340.000 55.000 39.977 −81.331 0.000 −2.699 2.349 −6.870 131.513 0.000 −2.236 −51.431 −3.031 −97.008 1.706 KP3_Ty1-G59Y_unrelaxed_rank_001_alphafold2_multimer_v3_model_3_seed_000_0001

##### Workflow design

You are working on a research project to use machine learning to develop antibodies or nanobodies for the newest variant of the SARS-CoV-2 spike protein that also, ideally, have activity against other circulating minor variants and past variants. Your team previous decided to modify existing nanobodies to improve their binding to the newest variant of the SARS-CoV-2 spike protein. Your team has built three components of a nanobody design pipeline: ESM, AlphaFold-Multimer, and Rosetta. Each of these three tools can be used to score a nanobody mutation based on how the mutation affects binding to an antigen. Your goal is to start with an existing nanobody and iteratively add mutations to it to improve its binding to the newest variant of the SARS-CoV-2 spike protein, resulting in 24 modified nanobodies with one or more mutations. Please determine how to use ESM, AlphaFold-Multimer, and Rosetta in this iterative design process. An important constraint is that ESM can evaluate all potential mutations to a nanobody in 5 minutes while AlphaFold-Multimer takes 30 minutes per mutation and Rosetta takes five minutes per mutation. The whole iterative process should take no more than a few days to complete. Note that AlphaFold-Multimer must be run before Rosetta on each mutation since Rosetta requires the nanobody-antigen structure predicted by AlphaFold-Multimer. Additionally, note that ESM log-likelihood ratios are generally in the range of 5-10 (higher is better), AlphaFold-Multimer interface pLDDT confidence scores are generally in the range of 60-80 (higher is better), and Rosetta binding energy scores are generally in the range of −20 to −40 (lower is better).

1. In each iteration, what is the order of operations for evaluating mutations with ESM, AlphaFold-Multimer, and Rosetta?
2. In each iteration, how many mutations (give a single number) will you evaluate with ESM, AlphaFold-Multimer, and Rosetta?
3. At the end of each iteration, how will you weigh the ESM, AlphaFold-Multimer, and/or Rosetta scores (give a formula) to rank the nanobody mutations?
4. At the end of each iteration, how many of the top-ranked mutations (give a single number) will you keep for the next round?
5. How will you decide how many iterations of mutations to run?
6. After all of the iterations are complete, how exactly (step-by-step) will you select the final set of 24 modified nanobodies from across the iterations for experimental validation?

##### Nanobody experimental validation

Codon optimized DNA sequences for the SARS-CoV-2 spike RBDs JN.1, KP.3, KP.2.3 and BA.2^26,53^, modified to include a N-terminal signal peptide (MFVFLVLLPLVSSQ), a C-terminal 6x his tag and a stop codon, were synthesized and cloned into pTwist-CMV-BetaGlobin (Twist Biosciences). RBDs were transiently expressed in Expi293 cells, and purified in parallel^54^ by Ni-NTA Excel affinity chromatography followed by desalting into PBS and concentration. The purification of Wuhan SARS-CoV-2 RBD has been described previously^54^. Codon optimized DNA sequences for nanobodies, modified to include a N-terminal pelB signal peptide (MKYLLPTAAAGLLLLAAQPAMA), a C-terminal 6x his tag and a stop codon, were synthesized and cloned into pET-29b(+) (Twist Biosciences). Nanobodies were expressed in 96-well and 24-well format in auto-induction media^55^, and periplasmic fractions from 4 mL of cell culture pellet were released by mild lysis in 400 uL PBS, following methods as described^56^.

Multiplexed ELISA measurements were performed as generally described^57^, with the following modifications. Two array patterns were printed using a sciFLEXARRAYER S12. The first pattern (Pattern 1) prints each spot in duplicate using a single 200-250uL drop for each spot. RBD antigens, BSA (negative control), and unmodified anti-Alpaca IgG VHH antibody (for total nanobody measurement) were printed at a source concentration of 50 ug/mL. The second pattern (Pattern 2), designed to increase the sensitive of binding, increased the number of drops per spot to 3, added a higher purity JN.1 RBD, and substituted the anti-Alpaca IgG VHH antibody with an anti-His IgG (for total nanobody measurement). Periplasmic fractions were diluted in PBS-T (5% skim milk in PBS + 0.05% Tween-20), and RBD-bound nanobodies were recognized by anti-Alpaca IgG VHH secondary antibodies (Jackson ImmunoResearch, 128-065-230 (for H11-D4, Nb21 and VHH-72 series), and 128-065-232 (for Ty1 series) at 1:10000 dilution in PBS-T.

